# DINOCT: robust skin surface localization for robotic noncontact dynamic optical coherence elastography

**DOI:** 10.64898/2026.09.05.749536

**Authors:** Raveeroj Bawornkitchaikul, Ruikang K. Wang, Matthew O’donnell, Ivan Pelivanov

## Abstract

Noncontact dynamic optical coherence elastography (OCE) can measure the elastic properties of anterior soft tissues. However, applying OCE to skin requires tracking mechanical waves propagating in multiple directions to reconstruct mechanical anisotropy. Robotic scanning is well suited to this task but requires accurate probe positioning, alignment to the surface normal, and precise, repeatable rotation. To ensure phase and polarization stability, polarization-maintaining (PM) fibers can be used, although they introduce stripe and ghost artifacts that can destabilize surface detection. We present DINOCT, a DINO-based method to localize the skin surface from optical coherence tomography (OCT) images that combines self-supervised pretraining with DINO (self-DIstillation with NO labels) and iBOT (image BERT pre-training with Online Tokenizer) objectives, low-rank adaptation, and a lightweight curve decoder. Compared with classical and supervised learning baselines, DINOCT achieved competitive localization accuracy on clean test data with a near-zero spike rate. Among the learned methods, DINOCT achieved the lowest average and maximum values of mean absolute error (MAE) for severe synthetic stress conditions and the lowest catastrophic failure rates on a separate recording with strong PM-fiber artifacts.

## 1. INTRODUCTION

Accurate localization of the tissue surface is essential for optical coherence tomography (OCT)-based applications that require precise probe positioning or are sensitive to sample motion during image acquisition. One such application is dynamic optical coherence elastography (OCE) of skin [1]. In dynamic OCE, mechanical waves are generated by an external source, such as a mechanical vibrator, a contact focused ultrasound (US) transducer, an air puff, or an air-coupled acoustic micro-tapping (A*µ*T) transducer [2, 3].

Mechanical wave propagation is tracked using phase-sensitive OCT by measuring wave-induced phase changes in the OCT signal, enabling reconstruction of mechanical wave fields and estimation of tissue elastic moduli.

Skin is mechanically anisotropic because of the local orientation of collagen fibers. Even in the simplest situation, where the local collagen fiber orientation can be approximated as unidirectional, the stiffness matrix **C** is defined by three independent elastic moduli, *G, µ*, and *δ* [1]; this differs from isotropic tissue, where the stiffness is determined by a single shear modulus *µ*.

The shear modulus of an isotropic, nearly incompressible medium can be reconstructed by measuring the velocity of shear or surface waves along an arbitrary direction inside the tissue volume or along its surface, respectively. Unfortunately, reconstructing elastic moduli in skin is not as simple and requires measurement of shear or Rayleigh wave propagation in different directions using phase-sensitive OCT [1, 4]. We previously showed that all three elastic moduli in skin can be reconstructed from Rayleigh wave speed angular anisotropy [1]. However, translating OCE into clinical practice requires precise control of the OCE scan-head position above the skin surface and precise rotation of the head to measure the angular dependence of the Rayleigh wave phase velocity. A robust way to control head positioning is to use a robotic arm in conjunction with the OCT scanning system [5].

To track Rayleigh wave propagation in different directions, the OCE scan-head must be aligned relative to the sample surface so that the target remains in the focal zone of both the OCT beam and the A*µ*T transducer. Head position should be maintained during each single angle scan, precisely rotated to a different angular position, and repeated for multiple propagation angles over the skin surface (in the (*xy*) plane). Accurate absolute positioning is essential to reconstruct spatially resolved elastic moduli. In addition, subject motion can introduce artifacts that can be compensated using co-registered structural OCT when robot positioning is well controlled [5, 6]. Errors in probe-to-surface distance or orientation can therefore affect acquisition consistency and the subsequent reconstruction of mechanical properties [5, 6].

Note that different OCT modalities have been used for structural and functional imaging in skin for a few decades [7–9]. For example, structural OCT is now widely used to identify skin tumor margins and assess tumor depth before Mohs micrographic surgery [10–12]. It also can evaluate burn depth and scarring [13– 15]. In aesthetic dermatology, OCT can be used to assess skin layers [16, 17]. Polarization-sensitive (PS) OCT can characterize and quantify collagen fiber orientation and even density, which, for instance, can be very useful in evaluating hypertrophic and atrophic scars [18–23]. OCT angiography (OCTa) is also used in dermatology to assess burn staging and surgery, characterize connective tissue disease, and evaluate cosmetic interventions [7, 14, 24, 25].

OCE is a relatively new modality, and it can be clinically useful if it can complement structural OCT, PS-OCT and OCTa. Indeed, the OCT/OCE system described in this paper is multi-modal and supports four modalities: structural OCT, PS-OCT, OCTa, and OCE. However, an integrated system can be very challenging. In particular, there is no consensus in the literature about using polarization-maintaining (PM) versus single mode (SM) fibers in polarization-sensitive (PS-OCT) and phase-sensitive (OCTa and OCE) measurements [26, 27].

SM-fiber OCT generally produces fewer structured artifacts but is highly sensitive to environmental vibrations, which can substantially change phase and polarization. This is especially critical for OCE, in which the displacement introduced by mechanical wave propagation may be on the order of 1/100 of the optical wavelength. Most SM-fiber systems require careful fiber attachment to a vibration-isolated breadboard. The OCT/OCE head in our system, however, undergoes fast robot motion. Under these conditions, achieving the required phase and polarization stability with freely moving SM fibers is not practical.

Although PM fibers can greatly stabilize OCT phase and polarization, they present a challenge for surface localization. PM-fiber OCT can produce structured artifacts, including horizontal coherence stripes associated with Fresnel reflections and vertically offset ghost copies caused by cross-coupling between orthogonal polarization modes [28]. In practice, these artifacts can create high contrast structures above or near the true skin surface and can destabilize classical gradient-based methods for surface detection and head alignment. The proposed DINOCT method, whose name combines DINO (self-DIstillation with NO labels) and OCT, is designed to reduce these artifact-induced surface localization failures while preserving the phase and polarization stability required for robotic multimodal OCT/OCE [6].

Thus, the goal of this paper is to develop and evaluate robust localization of the skin surface in OCT images used for robotic A*µ*T-OCE head positioning when there are significant artifacts introduced by PM fibers. We compare DINOCT with classical gradient methods and other learned methods.

Prior OCT studies in skin have primarily addressed epidermal segmentation, dermal-epidermal junction (DEJ) localization, and thickness estimation in bench top or offline settings using classical image processing or models based on convolutional neural networks (CNNs) [29–32]. Adjacent OCT surface segmentation work in other anatomies has focused on structured inference for retinal and corneal interfaces [33, 34].

Self-supervised pretraining in limited data visual domains is sensitive to the pretraining objective, architecture, training schedule, and target domain [35, 36]. Denoising-based pretraining has also been investigated for 3-D retinal OCT segmentation [37].

To our knowledge, prior work did not address these factors jointly for direct skin surface localization. The main contributions of this work are: (i) formulation of skin surface localization in robot-mounted PM-fiber OCT as direct centerline prediction in image coordinates; (ii) introduction of DINOCT, which combines self-supervised pretraining, parameter-efficient supervised adaptation, and a lightweight curve decoder; (iii) evaluation of DINOCT on relatively clean OCT images, severe synthetic corruptions, and a separate recording with strong PM-fiber artifacts; and (iv) comparison of DINOCT with competitive methods.

## 2. METHODS

### A. Multimodal A*µ*T-OCE imaging system

The multimodal A*µ*T-OCE imaging system combined an OCT scan-head and an air-coupled A*µ*T transducer mounted on a robot end effector. The robot-mounted, two-channel swept-source OCT system was designed to simultaneously collect received light with orthogonal polarizations to reconstruct polarization-sensitive OCT images in skin. In particular, local orientation of the optic axis was needed to define the orientation of collagen fibers in dermis independently from elastography measurements. OCTa can also be part of the scanning protocol. Thus, the platform supports structural OCT, PS-OCT, OCTa, and A*µ*T-OCE. Due to the PM-fiber OCT design, the circular polarization on the skin surface remained stable during all robotic manipulations and measurements and did not require additional adjustments after months of operation.

The OCT scan-head consisted of a fiber collimator, an XY galvanometer set (Cambridge Technology MicroMax™ Series 673, CT 6200H), and a scanning lens (LSM03, Thorlabs, USA). The PM-fiber arrangement used for polarization-sensitive and phase-sensitive OCT operation is shown in Fig. 1. The robotic arm hardware and scan geometry are shown in Fig. 2.

**Fig. 1.**
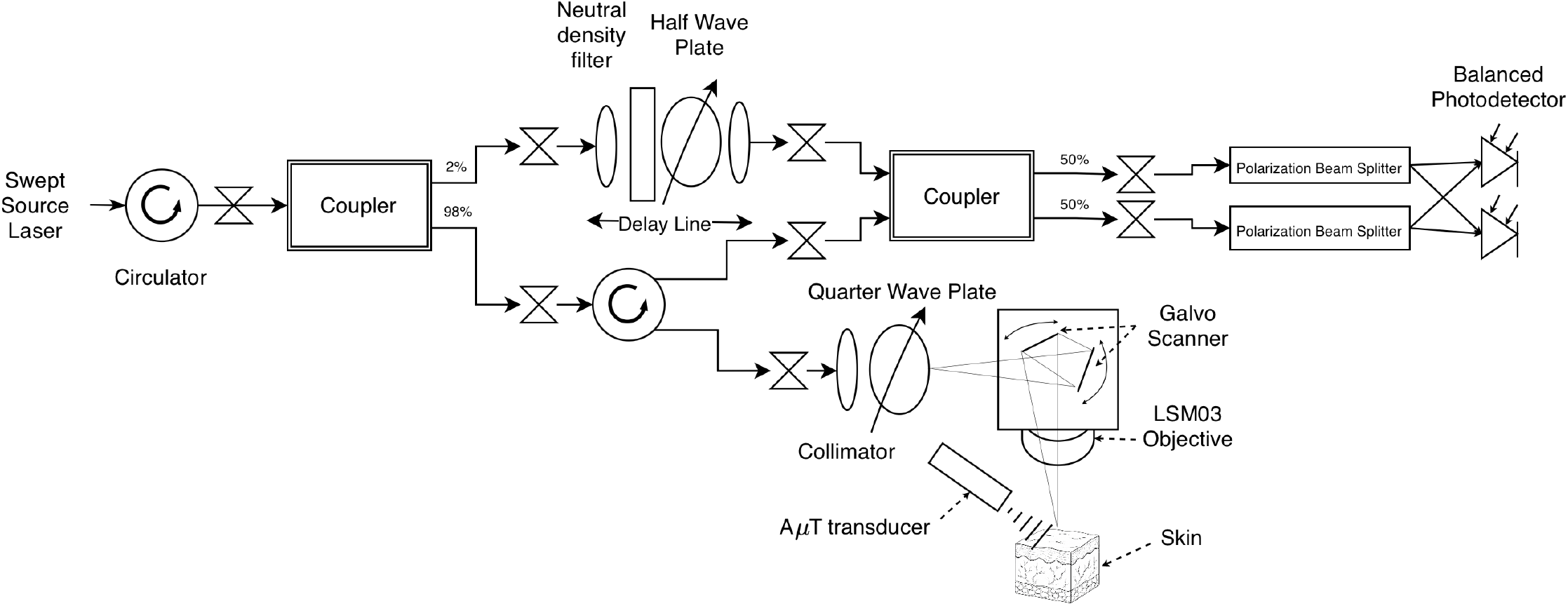
A simplified diagram of the noncontact, PM-fiber based, multimodal OCT/OCE system. All fiber components are made of **P**olarization-maintaining **AND A**bsorption-reducing (PANDA) style fibers.

**Fig. 2.**
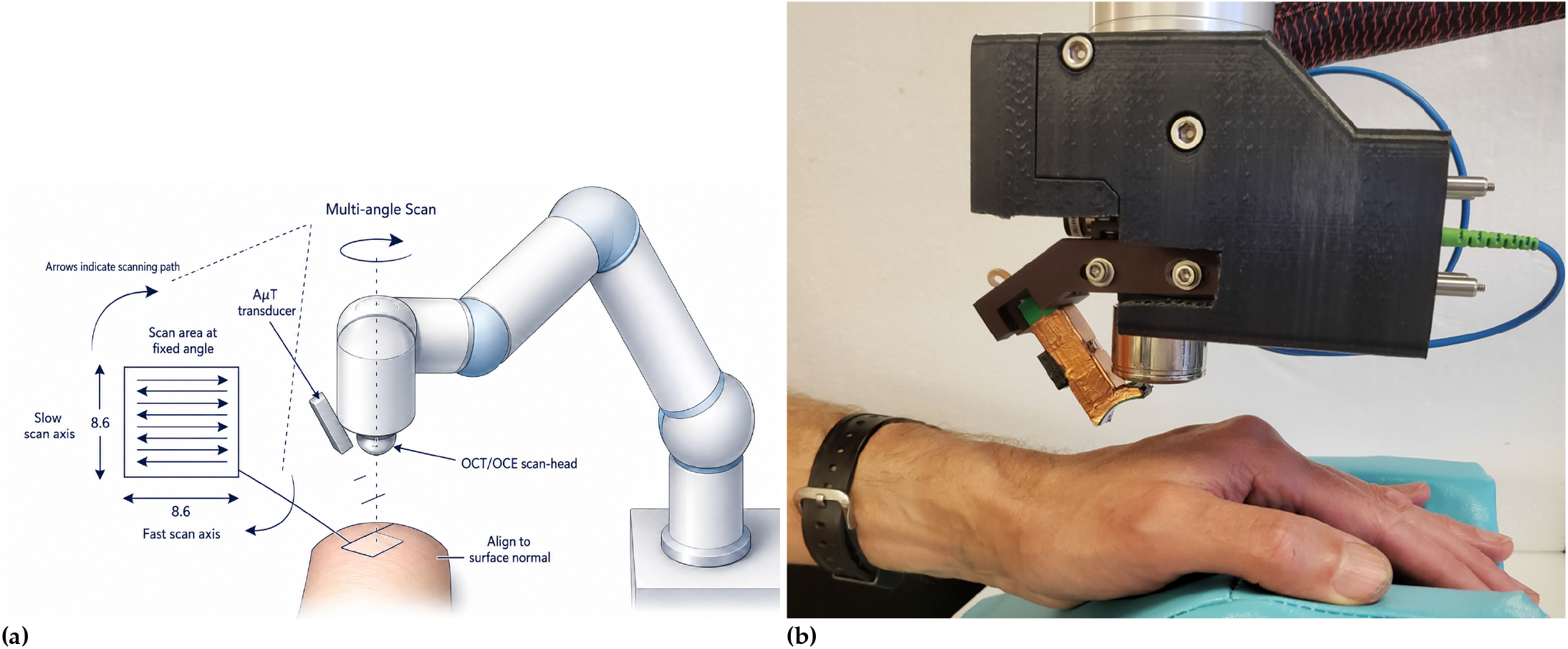
Schematics of noncontact robotic OCE. (a) Illustration of robot movement during scanning. The zoomed-in diagram on the left shows the OCT scanning path as it sweeps across the 8.6 × 8.6 mm surface area in a single 3D OCT scan. The end effector rotates about the estimated local surface normal to acquire scans at fixed angular positions spanning 180°. (b) Photo of end effector with the OCT head and A*µ*T transducer mounted on it, positioned above the human skin.

To generate elastic waves without contacting the skin surface, the cylindrically focused air-coupled ultrasound (A*µ*T) transducer applied a localized radiation force at the tissue surface, as described in our prior work [1, 3]. The measured noise displacement in the frequency range 200 Hz–2 kHz (the range of recorded Rayleigh wave signals in skin) was on the order of 20 nm, which was mainly defined by phase instabilities of the swept-source laser (Thorlabs, USA). Additional phase noise related to robotic mounting and the motion of the robotic arm was negligible. The position of the A*µ*T transducer was fixed during the OCE scan at a specified rotation angle. The signals from both photodetectors (PDB480C-AC, Thorlabs Inc., USA) were digitized with an ADC card (ATS9373, AlazarTech, Canada) at 1.8 GS/s for each channel.

DINOCT used structural OCT images to localize the skin surface (see Section B). The predicted surface was used to compute the probe-to-surface distance and define the local surface orientation. This information was used to autofocus the scan-head, maintaining it approximately perpendicular to the tissue surface at a specified axial offset [5]. Because the OCT head and A*µ*T transducer were rigidly mounted on the same end effector, aligning the OCT head automatically aligned the transducer. Optimal alignment produced a narrow acoustic line source, enabling the focused A*µ*T transducer to maximize both the displacement of the generated Rayleigh wave and its excitation bandwidth [3]. The positioning of the A*µ*T transducer relative to the OCT scanning lens is shown in Fig. 2(b).

Note that the present study focused on the problems of fast and robust skin surface localization, autofocusing and scan-head alignment. Structural OCT preview B-scans were inputs to DINOCT. The other modalities were not input to DINOCT. They were briefly introduced above for completeness; a detailed description of all modalities is outside the scope of this study.

### B. DINOCT architecture

DINOCT was trained in two stages. First, a ConvNeXt-Tiny backbone learned OCT image features from unlabeled B-scans using self-supervised learning (SSL). Second, low-rank adaptation (LoRA) modules and a lightweight curve decoder were trained with manually annotated surface centerlines while most backbone weights remained frozen. The model outputs one smooth skin depth profile 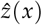 rather than a dense semantic segmentation map.

The backbone uses the standard ConvNeXt-Tiny scale, with stage depths (3, 3, 9, 3) and channel dimensions (96, 192, 384, 768). SSL pretraining used the multi-crop DINO self-distillation and iBOT (Image Bidirectional Encoder Representations from Transformers (BERT) Pre-training with Online Tokenizer), a masked-image objective with an online tokenizer, together with a Kozachenko–Leonenko (KoLeo) regularizer [39–42].

During post training, LoRA [43] modules with rank *r* = 8, *α* = 16, and dropout 0.02 were inserted only into `pwconv1`/`pwconv2` of the final six ConvNeXt blocks, with backbone normalization parameters also unfrozen. The component and LoRA placement sensitivity analyses supporting this selection are reported in Appendix, Section D (Tables 12 and 14). LoRA updates the original model weights *W* by training two low-rank decomposition matrices *A* and *B* in parallel such that an update is 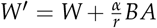, where 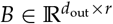 and 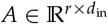 . Dropout *D*(*x*) is applied to the input during training, where the forward pass is 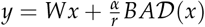.

The curve head is the task-specific decoder attached to the pretrained backbone. It first projects backbone features to 128 channels. Two depthwise 5 × 1 vertical convolutions aggregate evidence along the column direction and a residual depthwise 1 × 9 horizontal convolution couples neighboring columns to encourage local continuity of the predicted surface.

The curve-head design is anisotropic, because the desired output is one depth per column. Depth is where the evidence lies: the surface is an intensity step, so a vertical filter acts as an edge detector. Lateral is where the prior lies: the surface is continuous across columns, so the horizontal filter acts as a lateral continuity (smoothness) operator. Isotropic kernels with *k* × *k* convolutions could in principle express this, but it would conflate the evidence with the prior and is far more computationally costly.

Two stacked 5 × 1 convolutions give a 9-cell depth receptive field at nearly the same parameter cost as a single 9 × 1 layer, while the intervening nonlinearity adds expressiveness along the depth axis. The 1 × 9 horizontal convolution is zero-initialized with a learned per-channel scale, so the head begins as a pure per-column detector and training decides how much lateral coupling to introduce. Kernel sizes were selected empirically on validation data.

Because the head predicts a single depth per column, it is a constrained curve decoder rather than a segmentation decoder: it cannot represent arbitrary masks or multiple disconnected boundaries. The horizontal convolution further discourages sharp lateral surface transitions. Together these reduce the tendency to follow spurious high-contrast PM-fiber artifacts, such as stripes or ghost copies. The decoder was intentionally kept simple; exploratory multiscale, feature fusion, hybrid output, and expanded loss variants did not improve validation performance and were not retained.

The SSL pretraining configuration is summarized in Table 1, and the downstream post training configuration is summarized in Table 2 and shown in Fig. 3.

**Table 1.**
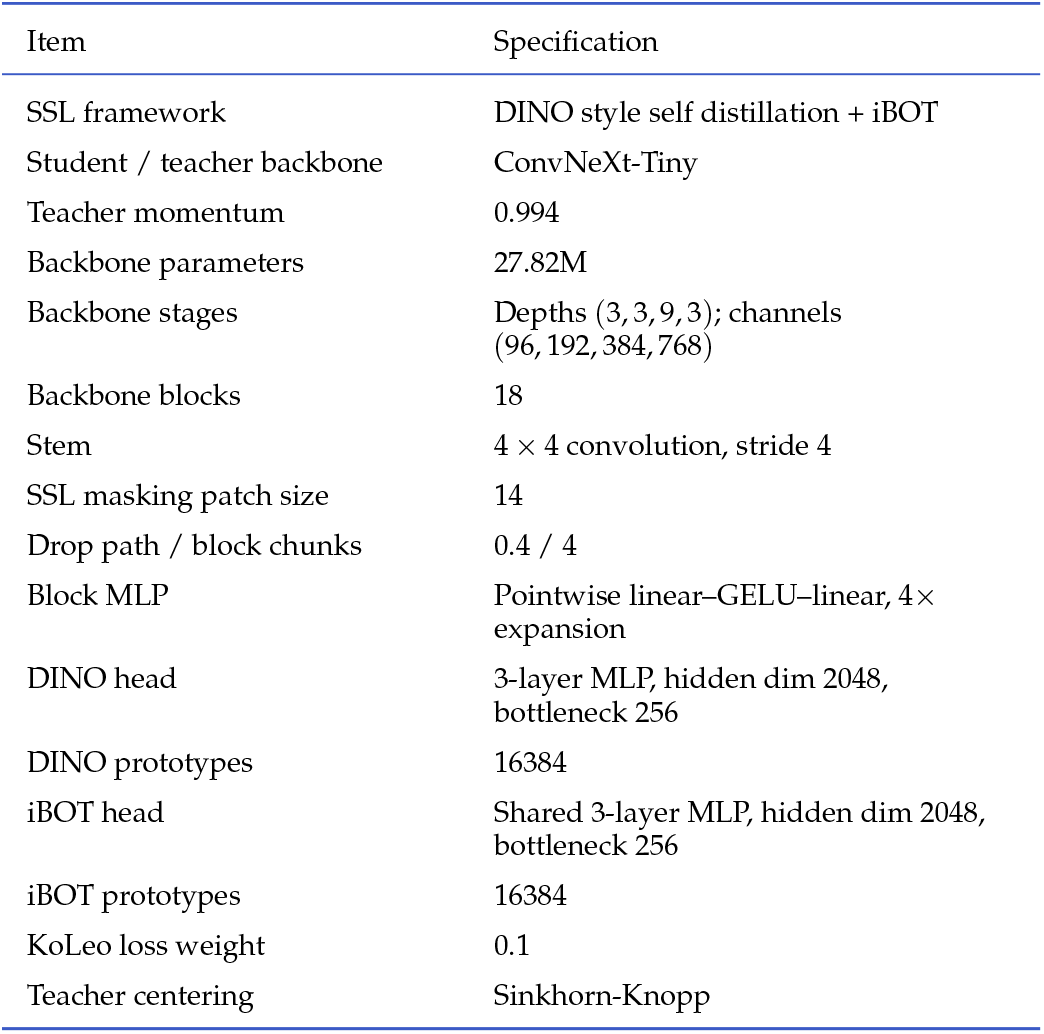
SSL pre-training configuration [45].

| Item | Specification |
| --- | --- |
| SSL framework | DINO style self distillation + iBOT |
| Student / teacher backbone | ConvNeXt-Tiny |
| Teacher momentum | 0.994 |
| Backbone parameters | 27.82M |
| Backbone stages | Depths (3, 3, 9, 3); channels (96, 192, 384, 768) |
| Backbone blocks | 18 |
| Stem | $4 \times 4$ convolution, stride 4 |
| SSL masking patch size | 14 |
| Drop path / block chunks | 0.4 / 4 |
| Block MLP | Pointwise linear-GELU-linear, $4 \times$ expansion |
| DINO head | 3-layer MLP, hidden dim 2048, bottleneck 256 |
| DINO prototypes | 16384 |
| iBOT head | Shared 3-layer MLP, hidden dim 2048, bottleneck 256 |
| iBOT prototypes | 16384 |
| KoLeo loss weight | 0.1 |
| Teacher centering | Sinkhorn-Knopp |

**Table 2.**
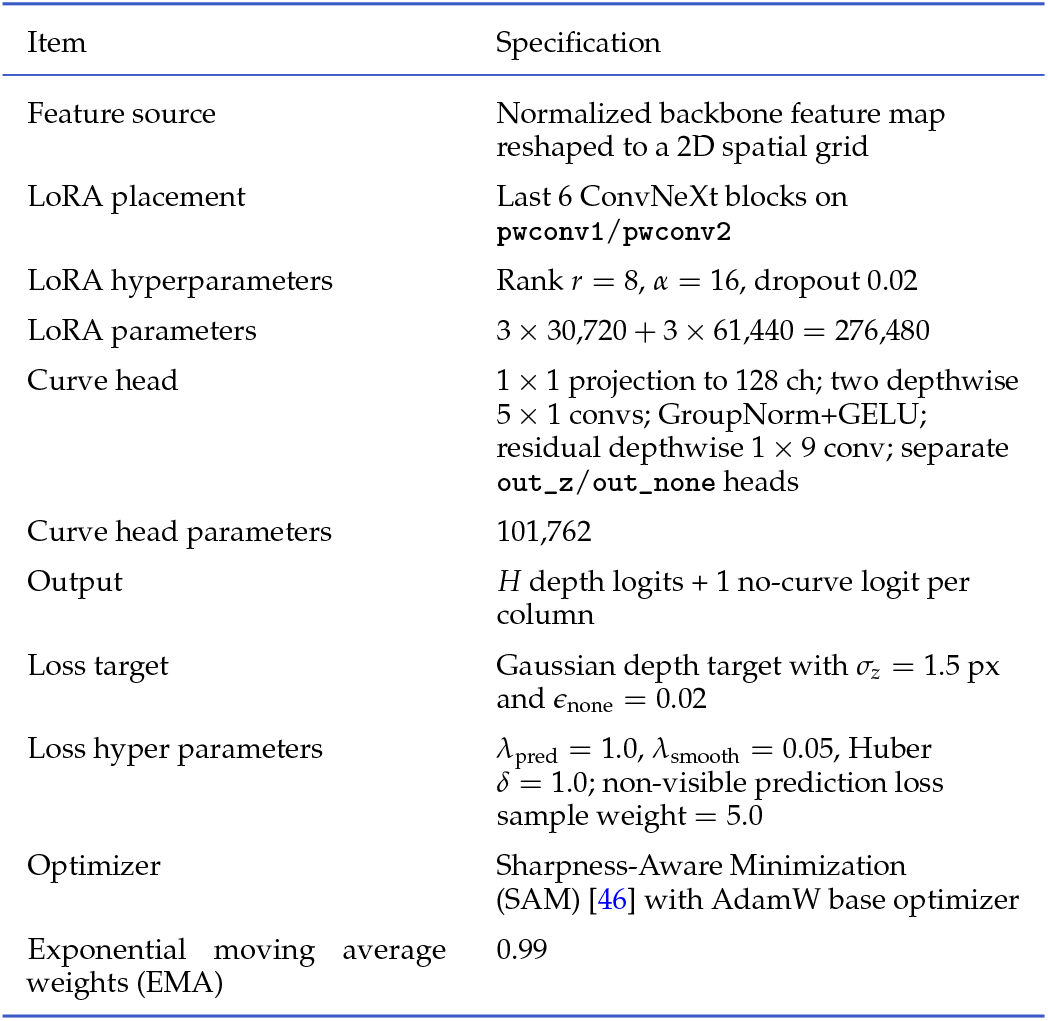
Post-training configuration [45].

| Item | Specification |
| --- | --- |
| Feature source | Normalized backbone feature map reshaped to a 2D spatial grid |
| LoRA placement | Last 6 ConvNeXt blocks on pwconv1/pwconv2 |
| LoRA hyperparameters | Rank $r = 8$, $\alpha = 16$, dropout 0.02 |
| LoRA parameters | $3 \times 30,720 + 3 \times 61,440 = 276,480$ |
| Curve head | $1 \times 1$ projection to 128 ch; two depthwise $5 \times 1$ convs; GroupNorm+GELU; residual depthwise $1 \times 9$ conv; separate out_z/out_none heads |
| Curve head parameters | 101,762 |
| Output | $H$ depth logits + 1 no-curve logit per column |
| Loss target | Gaussian depth target with $\sigma_z = 1.5$ px and $\epsilon_{\text{none}} = 0.02$ |
| Loss hyper parameters | $\lambda_{\text{pred}} = 1.0$, $\lambda_{\text{smooth}} = 0.05$, Huber $\delta = 1.0$; non-visible prediction loss sample weight = 5.0 |
| Optimizer | Sharpness-Aware Minimization (SAM) [46] with AdamW base optimizer |
| Exponential moving average weights (EMA) | 0.99 |

**Fig. 3.**
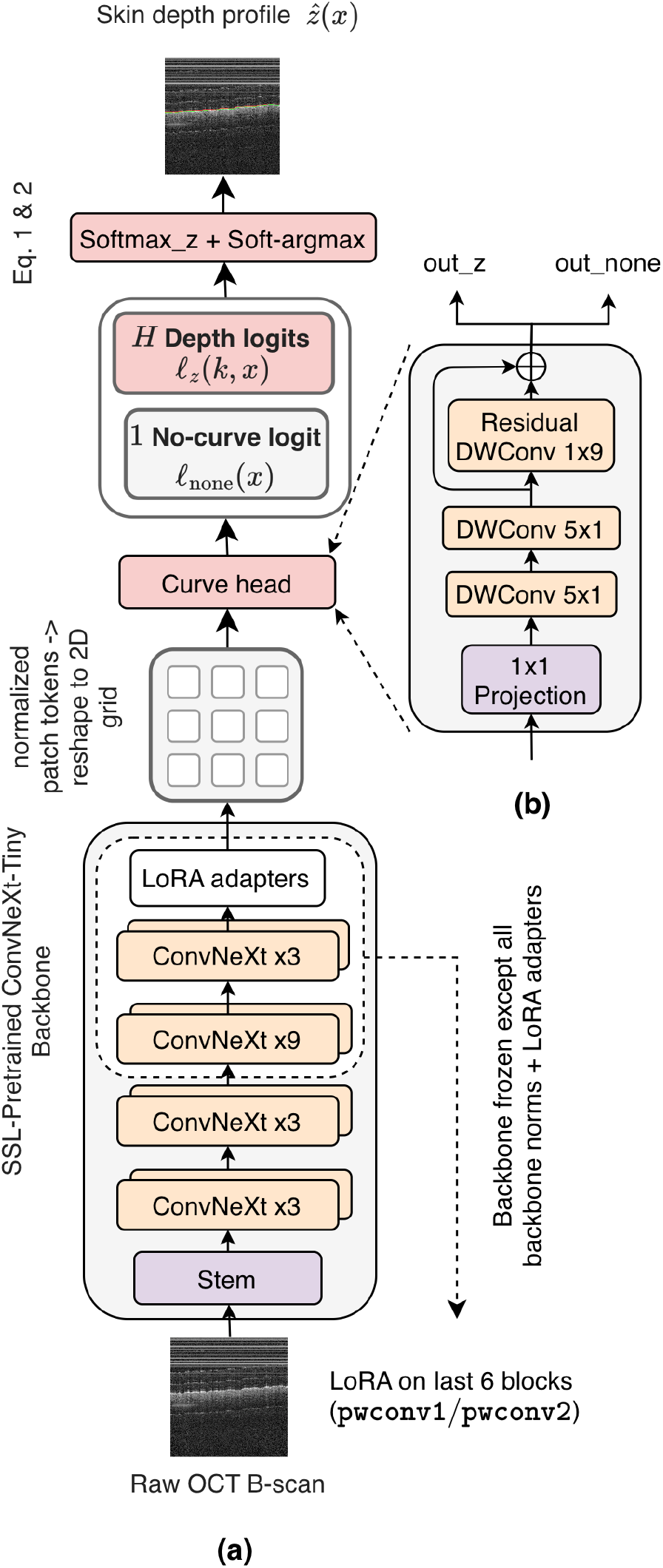
DINOCT architecture. (a) Overall pipeline for DINOCT. (b) Lightweight curve head. DWConv denotes depthwise convolution. ConvNeXt blocks follow [38]. Component and LoRA placement sensitivity analyses are provided in Appendix, Section D.

DINOCT used 500 SSL schedule epochs, with 1,250 optimizer iterations per epoch, using AdamW, with batch size of 32 per GPU, and a linear warm up followed by cosine learning-rate decay. Two 224-pixel global crops and eight 98-pixel local crops were used with color jitter, grayscale conversion, blur, solarization, and horizontal flips, following the DINO augmentation protocol. The student processed all crops, whereas the teacher processed only the two global crops. Crop sizes were kept as multiples of 14-pixel SSL masking patch size (Table 1). Because each image was normalized by its own mean and standard deviation, which is invariant to positive gain affine intensity changes, the brightness and contrast components of the color jitter had little effect on the network input, and the chroma components and grayscale conversion were exact no-op operations on these monochrome B-scans.

All augmentations were applied in the image domain before standardization. Standardization removes only positive-gain affine intensity changes, so the effects of blur, solarization, and geometric transforms persisted. Jitter, blur, and solarization strengths were increased relative to the reference configuration. In a matched comparison at equal training length, the increased setting substantially improved robustness to artifacts at comparable clean accuracy. Vertical flips were omitted in our setup and the reference configuration because they implied a different physical meaning.

Downstream post training used 1500 steps, corresponding to approximately 34 epochs over the 5,651 supervised post training frames (5,303 visible surface B-scans and 348 non-visible/background frames), batch size 128, Sharpness Aware Minimization (SAM) (*ρ* = 0.05) with an AdamW base optimizer, head/LoRA learning rates of 5 × 10^™4^ and 1 × 10^™4^, head/LoRA weight decays of 1 × 10^™3^ and 5 × 10^™5^, no data augmentation, and model selection by the lowest validation mean absolute error (MAE).

Given an input B-scan *I* ∈ R^*H*×*W*^, the backbone produced normalized patch features that were reshaped into a 2D spatial grid and passed to the curve head. With a 14-pixel patch size, a 512 × 500 B-scan is padded to 518 × 504 and encoded as a 37 × 36 token grid (the stage-4 ConvNeXt feature map, computed natively at stride 32, is bilinearly resampled to this grid), so the curve head operates at this token resolution; its output depth-logit map is bilinearly upsampled by a factor of 14 to full B-scan resolution before the per-column readout below. For each image column *x*, the curve head maps these features to *H* depth logits *l*_*z*_ (*k, x*), one for each possible axial depth *k* = 1, …., *H*, and one additional no-curve logit *l*_none_(*x*). A logit is an unnormalized score. The no-curve logit represents the alternative that no valid skin surface is visible at any of the candidate depths in that column. Together, the *H* depth logits and the no-curve logit form an *H* + 1-component logit vector for each column.

For each image column *x*, the depth logits *l*_*z*_ (*k, x*) were normalized across depth to form the predicted distribution of skin-surface depth:

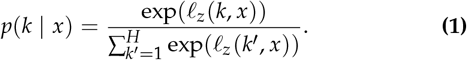

Here, *p*(*k* | *x*) represents the predicted probability that the skin surface is located at depth index *k* in column *x*. The predicted centerline is then computed as the expectation of this distribution:

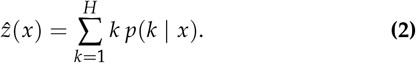

This expectation-based coordinate readout is related to differentiable spatial-to-numerical coordinate regression [44], applied here independently to each image column. Because the logit map is upsampled from the 37 × 36 token grid, sub-pixel localization arises from this expectation over the upsampled distribution rather than from token-level resolution. The resulting 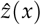 was the continuous-valued surface profile used for localization metrics and downstream surface fitting. The no-curve logit was used in the full *H* + 1-class training loss described below, but was not included in the depth-only soft-argmax used to recover the visible surface centerline.

The downstream DINOCT model contained 28.20 million parameters. During supervised post-training, the backbone was mostly frozen; only the curve head, LoRA adapters, and backbone normalization parameters were trainable. This resulted in 0.395 million trainable parameters, corresponding to 1.40% of the full model. Of these, 0.102 million parameters were in the curve head and 0.276 million were in the LoRA adapters, with the remaining 16,320 in trainable backbone normalization parameters.

### C. Training objective

For simplicity, the reduced notation of indexes was used as described below:

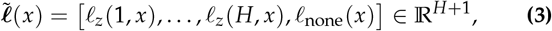

where the first *H* entries correspond to the estimated depth logits and the (*H* + 1)th entry corresponds to the no-curve logit. The no-curve class provided auxiliary supervision for non-visible scans and formed the basis of the surface presence criterion described below, which was used only during deployment. Standalone performance of this rejection criterion was not evaluated in this study.

The full prediction distribution is

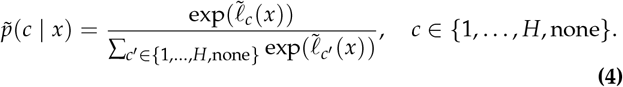

Accordingly, for depth classes *k* = 1, …., *H*,

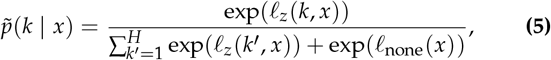

and for the no-curve class,

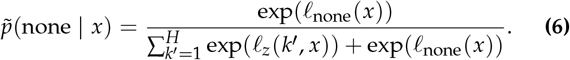

For deployment, the per-column no-curve probabilities were reduced to a surface presence score for each B-scan:

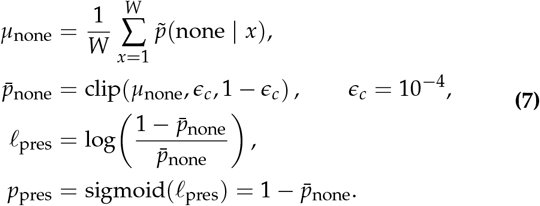

Here, sigmoid(·) denotes the logistic sigmoid function. The exported ONNX output *l*_pres_ is therefore a deterministic reduction of the per-column no-curve probabilities rather than a separately trained prediction head. The deployment threshold *p*_pres_ ≥ 0.70 is equivalent to 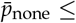 0.30. The value 0.70 was chosen heuristically as a deployment engineering criterion and was not optimized as part of the reported experiments.

The full distribution 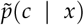 was used in the prediction loss, whereas the depth only distribution *p*(*k x*) was used for centerline recovery.

For scans with a valid visible skin surface, the depth target was constructed as a soft per-column label centered at the annotated surface depth *z*⋆(*x*). Specifically, we used a discrete Gaussian target over axial pixel index *k*, with standard deviation *σ*_*z*_ = 1.5 px:

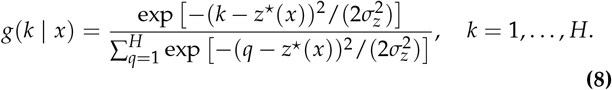

The value *σ*_*z*_ = 1.5 px defines the width of the soft training target around the manual centerline and was used only for supervised training. A small probability mass *ϵ*_none_ = 0.02 was assigned to the no-curve class. For scans without a valid visible skin surface, the full target mass was assigned to the no-curve class.

For visible-surface scans,

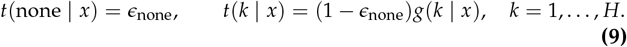

For non-visible scans,

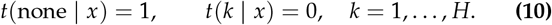

The prediction loss is the column-wise cross-entropy over the full (*H* + 1)-class distribution:

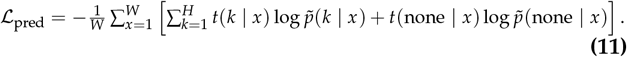

During post training, non-visible scans received a prediction-loss weight of 5.0 to partially compensate for their lower frequency (348 non-visible versus 5,303 visible training scans) and to maintain learning of the no-curve output. The value worked well empirically but was not formally tuned or included in a separate sensitivity analysis.

To suppress spikes while preserving valid local curvature, we added a smoothness term on the soft argmax centerline:

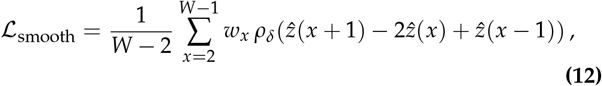

where *ρ*_*δ*_ (·) denotes the Huber penalty and *w*_*x*_ is an entropy derived confidence weight computed from the predicted depth distribution. The smoothness term was applied only to visible-surface training scans; non-visible scans contributed only to *ℒ*_pred_.

The confidence weight was defined as

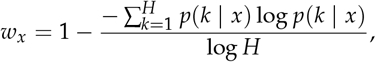

so that high entropy column predictions contributed less to the smoothness term.

The total loss was defined as

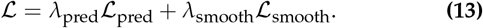

The coefficient *λ*_pred_ = 1 sets the reference scale of the prediction loss and *λ*_smooth_ = 0.05 sets the scale for local smoothing effects. After the horizontal convolution layer was adopted in the curve-head, the need for *λ*_smooth_ has been substantially reduced. Since we desire a more expressive skin profile, the current smoothness term is reduced to provide a very weak local smoothing effect; removing it produced no change at the reported precision in the component analysis. The coefficients were not jointly optimized.

### D. Acquisition protocol and 3D preview scans

The OCT/OCE scanning protocol included three consecutive steps: (i) OCT-based autofocusing and surface normal alignment before scanning, (ii) optional PS-OCT and OCTA acquisitions, and (iii) A*µ*T-OCE acquisition. During OCE acquisition, the robot rotated the OCT/OCE scan-head from 0° to 180° about the estimated local surface normal. The number of angular positions could be adjusted according to the requirements of the elastic moduli reconstruction. To compensate for possible subject motion, intermediate structural OCT preview scans and autofocus updates could be acquired between OCE angular positions.

The maximum OCT scanning area was 8.6 × 8.6 mm, although smaller rectangular regions could be used. For a centered 8.6 mm scan, the rotation trajectory for the selected scan point followed a semicircle with a 4.3 mm radius centered at the OCT scan center. For smaller or shifted scan regions, the rotation center could be displaced so the OCT scan remained close to the A*µ*T excitation line, where Rayleigh wave magnitude was highest.

For each sparse volumetric preview, six B-scans were recorded during the galvanometer slow-axis sweep, with each B-scan formed by a fast-axis sweep (Fig. 4). Together, these B-scans formed the sparse volumetric input to the surface-localization and point-cloud construction procedure described in Section E.

**Fig. 4.**
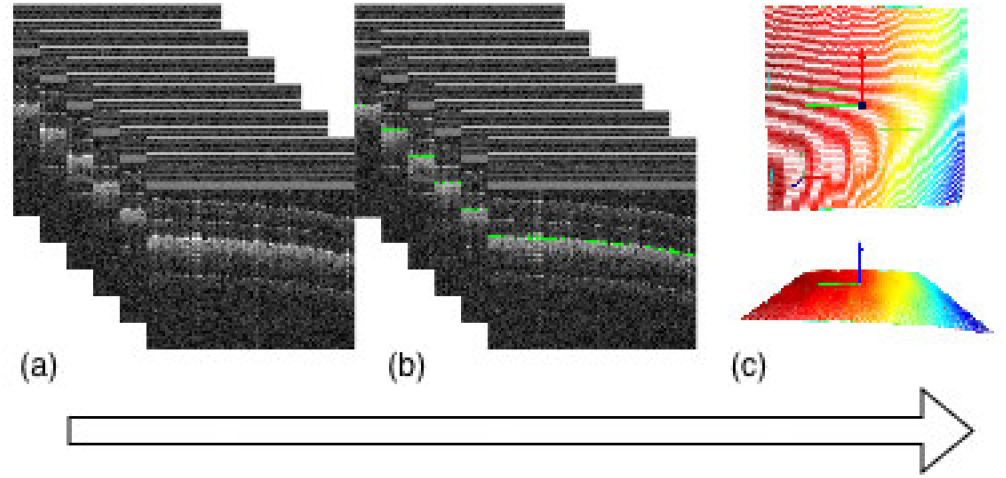
Surface localization pipeline used for OCT/OCE head positioning. The image bundle in panel (a) shows the sparse 3D OCT preview scan represented as a stack of B-scans. The surface localization algorithm estimates one candidate axial depth profile for each B-scan, shown as overlays in image bundle (b). Only profiles retained by the surface-presence criterion are converted into a sparse 3D point cloud by assigning each profile a uniformly spaced coordinate *sj* along the galvanometer slow scanning axis in the OCT frame; rejected B-scans contribute no points. Panel (c) shows top and isometric views of the detected sparse surface points and the fitted plane. The color gradient represents depth; red and blue indicate higher and lower values, respectively. The interpolated surface is shown only for visualization. In practice, plane fitting uses the sparse detected points. The fitted plane provides the centroid and rotation matrix used by the robot path planning layer. Only the axial component of the centroid is converted from pixels to a metric focus height correction using the calibrated pixel-to-mm scale factor.

### E. Surface-based OCT/OCE head positioning

Surface localization was used to convert sparse OCT preview scans into robot position updates for autofocusing and OCT/OCE head alignment. This study evaluates surface localization in image coordinates.

During an autofocus update, the system acquired a sparse structural OCT preview volume as a sequence of B-scans sampled during the sweep of the galvanometer slow axis. For the *j*th preview B-scan, the model produced two outputs: a candidate one-dimensional axial surface profile 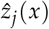 (*x* is the fast axis image coordinate and *z* is OCT depth) and a scalar surface-presence score indicating whether a usable surface was visible. The score was derived from the per-column no-curve probabilities as defined by Eq. (7). The exported model returned the corresponding logit *l*_pres,*j*_, with *p*_pres,*j*_ = sigmoid(*l*_pres,*j*_ ). The candidate profile was retained only when *p*_pres,*j*_ ≥ 0.70; otherwise, the B-scan contributed no surface coordinates to the reconstructed surface. The retained profiles were converted into a sparse 3D surface point cloud by assigning each profile an evenly spaced slow axis coordinate *s*_*j*_:

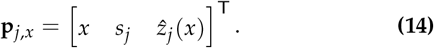

Thus, the preview stack was represented as sparse surface points in image coordinates. Robust plane fitting was attempted on these points only when at least two retained B-scans were available at distinct slow axis positions. If fewer than two B-scans were retained at distinct slow scanning axis positions, or if the robust plane fit failed, no pose update was produced from that preview. If autofocus remained enabled, the system automatically acquired another preview and repeated surface estimation. The fitted pose was subsequently transformed from the OCT frame to the robot frame using the calibrated OCT-to-robot coordinate transformation. Interpolated surfaces were used only for visualization.

Figure 4 summarizes the pipeline for surface localization and position update. The pipeline used a robust local plane fit as the primary surface estimator. The fitted plane was represented as

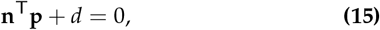

where **p** = [*x, s, z*]^T^ is a point in image coordinates, **n** is the estimated local surface normal, and *d* is the plane offset. The fit was computed using Tukey-weighted residuals to reduce the influence of isolated outlier columns and local spike-like detections:

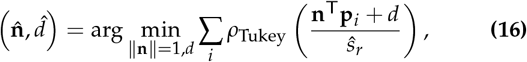

where **p**_*i*_ denotes a detected sparse surface point and *ŝ*_*r*_ is a robust residual scale estimate. We define *ŝ*_*r*_ = *k* MAD, where *k* is a constant scale factor and MAD is the median absolute deviation of the current point-to-plane residuals. For normally distributed residuals, *k* ≈ 1.4826. For the present system, the approximate axial calibration was 65 pixels/mm, or about 0.015 mm/px. This calibration was used only to convert the axial focus correction.

The robust fit returned a centroid 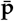 and a rotation matrix. For autofocus, only the axial component of the centroid was converted into a metric height correction:

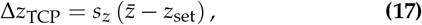

where Δ*z*_TCP_ is the commanded axial displacement of the robot tool center point (TCP), 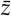 is the fitted centroid depth in pixels, *z*_set_ is the desired image space surface depth, and *s*_*z*_ is the calibrated axial scale factor in mm/px. The fitted rotation matrix was passed to the robot path planning layer to update OCT/OCE head orientation. The lateral components of the centroid were not used as independent lateral robot corrections during this autofocus step; lateral scan positioning was handled by the scan plan and robot trajectory definition.

Earlier versions of the positioning pipeline estimated the local orientation using a minimum volume, oriented bounding box fitted to detected surface points [47, 48]. However, this approach was sensitive to outliers because the box had to enclose the full point set. A single local spike-like surface localization error could therefore expand the box and rotate its estimated axes away from the expected local surface orientation, as shown in Fig. 5. Such a failure mode motivated replacing the bounding box orientation estimate with the current robust plane fitting procedure.

**Fig. 5.**
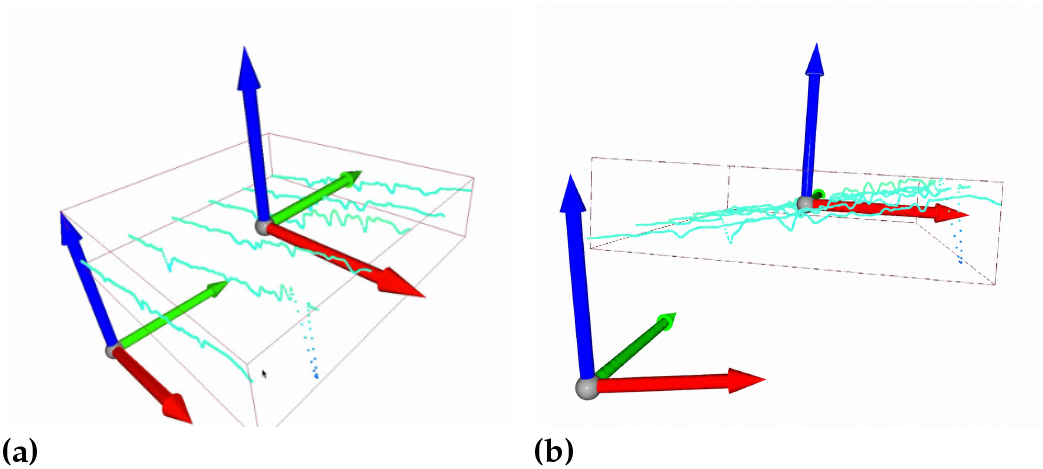
One of the failure cases for an earlier estimator based on the orientation of a minimum volume oriented bounding box. (a) Isometric view of the fitted point cloud and coordinate frame. (b) Side view showing the local downward shift caused by the surface localization outlier. The outlier expands the fitted bounding box [47, 48] downwards and rotates its estimated axes away from the expected local surface orientation.

Figure 6 visualizes the coordinate frame update for a successful OCT/OCE head positioning case.

**Fig. 6.**
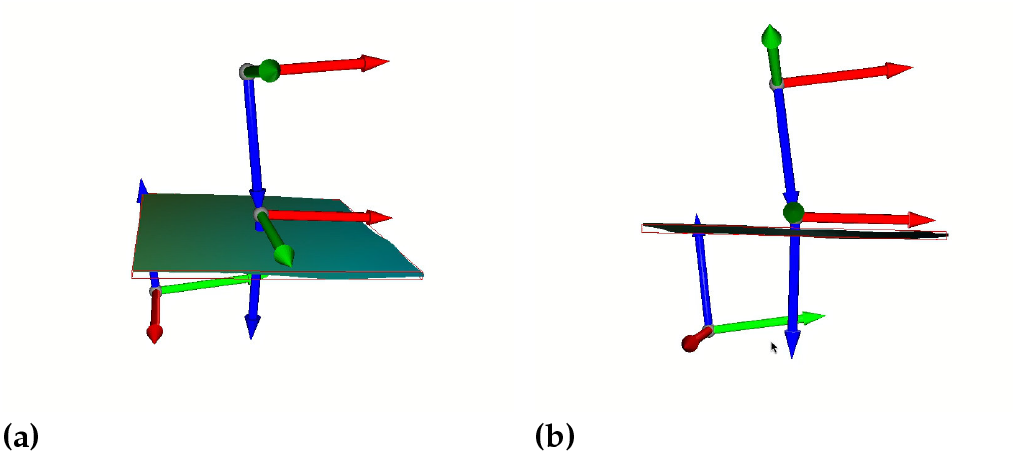
Coordinate frame visualization for a successful OCT/OCE head positioning update. (a) Isometric view showing three coordinate frames and interpolated surface. (b) Side view showing alignment of a coordinate frame on the interpolated surface. The upper coordinate frame denotes the unaligned robot/end effector frame, and the coordinate frame near the reconstructed surface denotes the robot/end effector frame after applying the fitted rotation matrix. The interpolated surface is shown only for visualization; plane fitting uses detected sparse surface points. The coordinate triads are qualitative renderings of the commanded frame update. The displayed axial axis should not be interpreted as a separately calibrated estimate of the surface normal.

### F. Human subjects

Healthy adult nonsmoking volunteers without known skin disease were scanned at multiple skin sites as part of a broader study of baseline skin mechanical properties. More than 30 subjects were included in the broader imaging study. All imaging procedures in human subjects were approved by the University of Washington Institutional Review Board (STUDY00012306), and written informed consent was obtained from all participants.

The present study used structural OCT data to develop and evaluate robotic surface localization and auto-focusing algorithms. Reconstruction of elastic moduli from OCE measurements was outside the scope of this study; related results can be found in our previous work [1, 4], and will be published separately.

Training, validation, and test assignments were made at the recording/acquisition level, so frames from the same continuous sequence or 3-D scan were not split across partitions. Participant and skin-site identifiers were not used for DINOCT with the goal of making it robust and stable across all possible image sites and healthy subjects.

### G. Data and annotations

Dataset composition and split accounting are summarized in Tables 3 and 4. The data included two structural OCT acquisition modes: continuous free motion B-scan sequences and B-scan slices extracted from sparse volumetric preview acquisitions.

**Table 3.** Visible surface labeled dataset summary.

| Subset | Acquisitions / recordings | Visible labeled B-scans | Split rule | Image size |
| --- | --- | --- | --- | --- |
| Continuous sequences | 18 | 536 | by sequence | 512 × 500 |
| B-scans derived from 3-D datasets | 1,462 | 6,810 | by C-scan | 512 × 500 |
| Stress recording | 1 | 151 | evaluation only | 512 × 500 |

Each raw OCT B-scan had dimensions *H* × *W* = 512 × 500 pixels. The horizontal image coordinate corresponds to the OCT fast axis direction, and the vertical image coordinate corresponds to OCT depth. Each visible surface label was a manually drawn centerline with one depth index per image column. Method-specific targets, such as segmentation bands or probability distributions, were used only during training; all reported localization metrics were computed against the same manual centerlines.

Training, validation, and testing assignments were made at the recording/acquisition level, so frames from the same 3D OCT scan were not split across sets. Non-visible/background frames did not contain a usable skin surface because the surface was out of view, dominated by a specular reflection, or insufficiently visible. The 348 training frames in this category were used for SSL pretraining and auxiliary no-curve supervision; the 150 validation/test non-visible frames were not used for training and were excluded from localization metrics (Table 4).

**Table 4.**
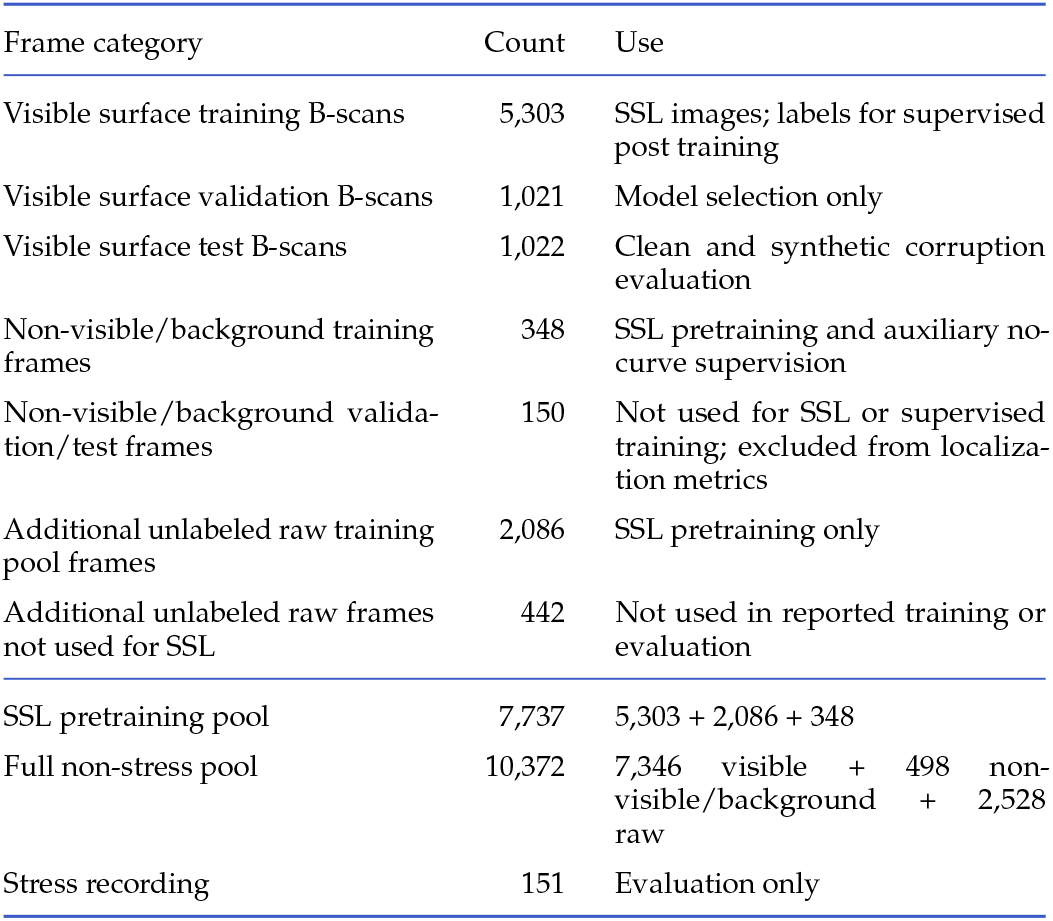
Split and SSL pretraining accounting. The stress recording was excluded from training and formal model selection.

SSL pretraining used only recordings assigned to training and did not use manual centerline labels. Validation and test recordings were excluded from SSL pretraining and supervised parameter updates.

We also evaluated 151 B-scans with visible skin surface from a separate recording acquired using a hardware configuration that produced unusually strong PM-fiber artifacts. This case-specific stress test is referred to below as the stress recording. It was excluded from training and formal model selection. Although its predictions were inspected during development, no checkpoint, architecture, optimizer, or hyperparameter choice was based on this recording.

### H. Comparison methods

We compared DINOCT against other deep learning (DL) supervised methods and classical gradient-based algorithms commonly used in OCT image segmentation and feature extraction [29, 30]. DL baselines were included to reflect prior CNN-based skin OCT analysis [31, 32] and adjacent OCT surface localization approaches [33]. All DL methods used the same labeled split for training/validation/testing and model selection. All DL methods were evaluated using the same pixel domain and centerline metrics. Accordingly, the main comparison evaluates complete surface-localization methods under matched data and evaluation conditions; it is not intended to isolate the effect of backbone architecture alone. Additional implementation details, including the GF-B and GRAD-SG variants and complete U-Net and FCBR-style architecture descriptions, are provided in Appendix, Section A.

#### GF

A minimal gradient filter baseline consisted of row-wise subtraction of the mean for DC suppression, Gaussian smoothing, positive vertical gradient scoring, and column-wise peak picking.

#### GRAD-ENG

Engineered gradient surface detection was based on row-wise DC suppression, low-pass filtering, intensity-weighted gradient scoring, median filtering, heuristic outlier removal, and Kalman smoothing (see Fig. 7).

**Fig. 7.**
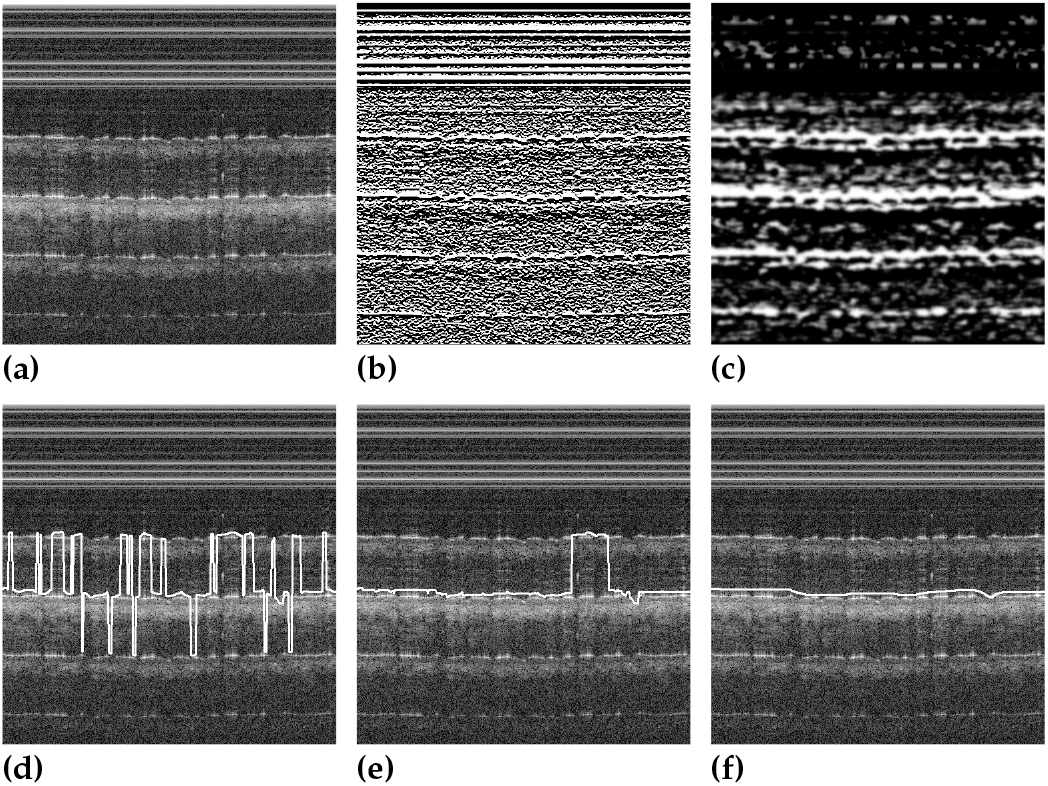
GRAD-ENG processing sequence. (a) Raw B-scan OCT image. (b) B-scan OCT image after gradient and low-pass filtration applied. (c) Image (b) after row-wise DC suppression. (d) Computation of column-wise maximum. (e) Outlier suppression. (f) Final smoothing and Kalman-filtered result. Additional classical-method implementation details are provided in Appendix, Section A.

#### TUNED-SOBEL-DC

This type of surface detection was implemented in an earlier bench pipeline. It was based on a vertical Sobel-style response, fixed row DC suppression, heavy spatial smoothing, and heuristic outlier cleanup.

#### U-Net (UNet)

A UNet-style encoder–decoder [49] was trained from scratch as a supervised surface-localization baseline. The model used four encoder stages, a bottleneck, and four decoder stages with skip connections. Its final feature map was passed to a curve distribution head that predicted *H* depth logits and one no-curve logit for each image column. The model was trained using the learned baseline protocol described in Appendix, Section A.1, and was evaluated using the same centerline metrics as the other methods.

#### FCBR-style boundary regressor (FCBR)

A lightweight fully convolutional boundary regressor was trained from scratch as a supervised surface-localization baseline. The model uses a shallow convolutional stem, depthwise separable downsampling blocks that preserved lateral resolution, and a dilated context stack. It was trained using the learned baseline protocol described in Appendix, Section A.1, and was evaluated using the same centerline metrics as the other methods. We report this model as an FCBR style re-implementation [50] rather than an exact historical reproduction. Full architecture and training details are provided in Appendix, Section A.1.

### I. Evaluation protocol and metrics

All metrics were computed against manually drawn reference centerlines in image coordinates. During training, methods were exposed to non-visible frames using their native supervision schemes, while reported localization metrics were computed only on visible surface scans. The surface presence threshold was not applied when computing visible-surface localization metrics; standalone performance of the rejection criterion was not evaluated. Robot positioning used calibrated OCT-to-robot scale factors. Localization accuracy is reported here in pixels to make it more convenient for implementation on different OCT systems independent of the depth sampling rate. This is consistent with recent skin OCT work, which reports boundary accuracy in pixels and notes that manual agreement is generally acceptable within two pixels [32].

To prevent leakage within the acquisition, train/validation/test splits were performed at the recording level: all frames from a continuous sequence were assigned to one split, and all slices from a 3-D acquisition were assigned to one split. We report mean absolute error, signed bias, tolerance-based accuracy, spike rate, and runtime. Error and quality metrics were computed per B-scan and averaged within each recording. For learned methods, the five seed-specific values were then averaged within each recording, and the reported mean ± standard deviation was computed across recordings. Classical baselines were deterministic single runs and were summarized across recordings. Runtime is reported separately as mean ±standard deviation over repeated per-B-scan timing measurements. For the stress recording, metrics were summarized across B-scans rather than across recordings; this exception is reported separately from the summaries for clean and synthetic corruption tests. After model selection, no method or hyperparameter was adjusted separately for an individual evaluation recording. Each method used a fixed inference procedure, and all predicted centerlines were scored using the same metric implementation.

Let 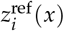 denote the manually drawn reference surface depth for test image *i* at A-scan *x*, and let 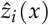 denote the predicted surface depth. For the network outputs of a depth distribution per column *p*_*i*_ (*k*|*x*), the predicted centerline was obtained using Eq. (2).

#### Mean absolute error in pixels

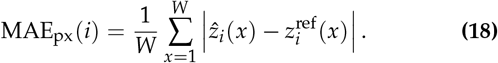

Eq. (18) averages the absolute column-wise depth error over all *W* lateral image columns in B-scan *i*.

#### Signed bias in pixels

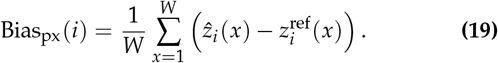

Eq. (19) averages the signed column-wise depth error. Because the OCT depth coordinate increases with image depth, a positive value indicates that the predicted surface is, on average, deeper than the manual reference, whereas a negative value indicates a shallower prediction.

#### Position accuracy at tolerance *τ*

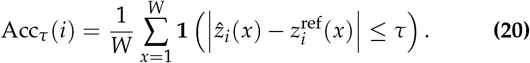

Eq. (20) reports the fraction of columns whose predicted surface depth is within *τ* pixels of the manual reference. In the result tables, Acc@2px corresponds to *τ* = 2 px and gives the fraction of A-scans with near-reference surface localization.

##### Spike / outlier rate

To quantify unstable local jumps in the predicted surface, we measured the fraction of columns whose second-order difference exceeded a threshold *κ*:

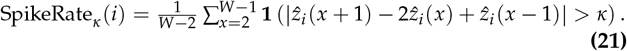

The threshold *κ* = 1.0 px was selected from the validation set based on the upper tail of second order differences observed in manual reference centerlines at the 0.99 quantile. In all result tables, SpikeRate_*κ*_ is reported in percent.

##### Catastrophic failure rates for B-scans

These rates summarize whether an entire preview frame would be considered unreliable for position updating. For the stress recording, we also report failure rates for individual B-scans. FailureRate@*T* is the percentage of B-scans whose MAE exceeds *T* pixels. The B-scan spike/failure rate is the percentage of B-scans containing at least one column whose second order difference exceeds the spike threshold *κ*. For learned methods, these B-scan failures and spike rates are computed per training seed and then averaged, so they reflect single model behavior rather than ensemble behavior. These metrics are intended to capture rare large localization failures that may be more relevant to robot position updates than the mean clean set error alone.

##### Runtime

We report average end-to-end wall clock latency per B-scan on the deployment hardware (ASUS NUC 14 Pro with Intel Core Ultra 5 125H, 16 GB RAM, 512 GB NVMe). Learned models were exported to Open Neural Network Exchange (ONNX [51]) format and evaluated on OpenVINO, whereas gradient-based baselines were timed in their native CPU Python implementations on the same machine. These runtime values reflect the current deployment implementations rather than language-independent algorithmic complexity.

### J. Artifact robustness protocol

To assess performance with significant image artifacts, we evaluated each method on the clean test images and on synthetic variants of the same images. The synthetic corruptions were not used for SSL pretraining or supervised parameter updates. Results for severe corruption were examined during method development and informed the choice of LoRA placement. We therefore treat this as a stress comparison used during development of the method rather than as an independent test.

The benchmark included horizontal stripes, vertically shifted ghost copies, and local regions with reduced contrast. Each corruption was evaluated at medium and severe levels. The operators were designed to resemble common artifacts in PM-fiber OCT, but not to reproduce their complete physical origin. Figure 8 shows representative examples. Stripe corruption added bright horizontal bands with a Gaussian intensity profile along the axial direction. Ghost corruption added a vertically shifted copy of the image. Contrast corruption added soft localized regions, with raised cosine edge profiles, of reduced intensity and contrast. Detailed operators and parameters are reported in Section B.

**Fig. 8.**
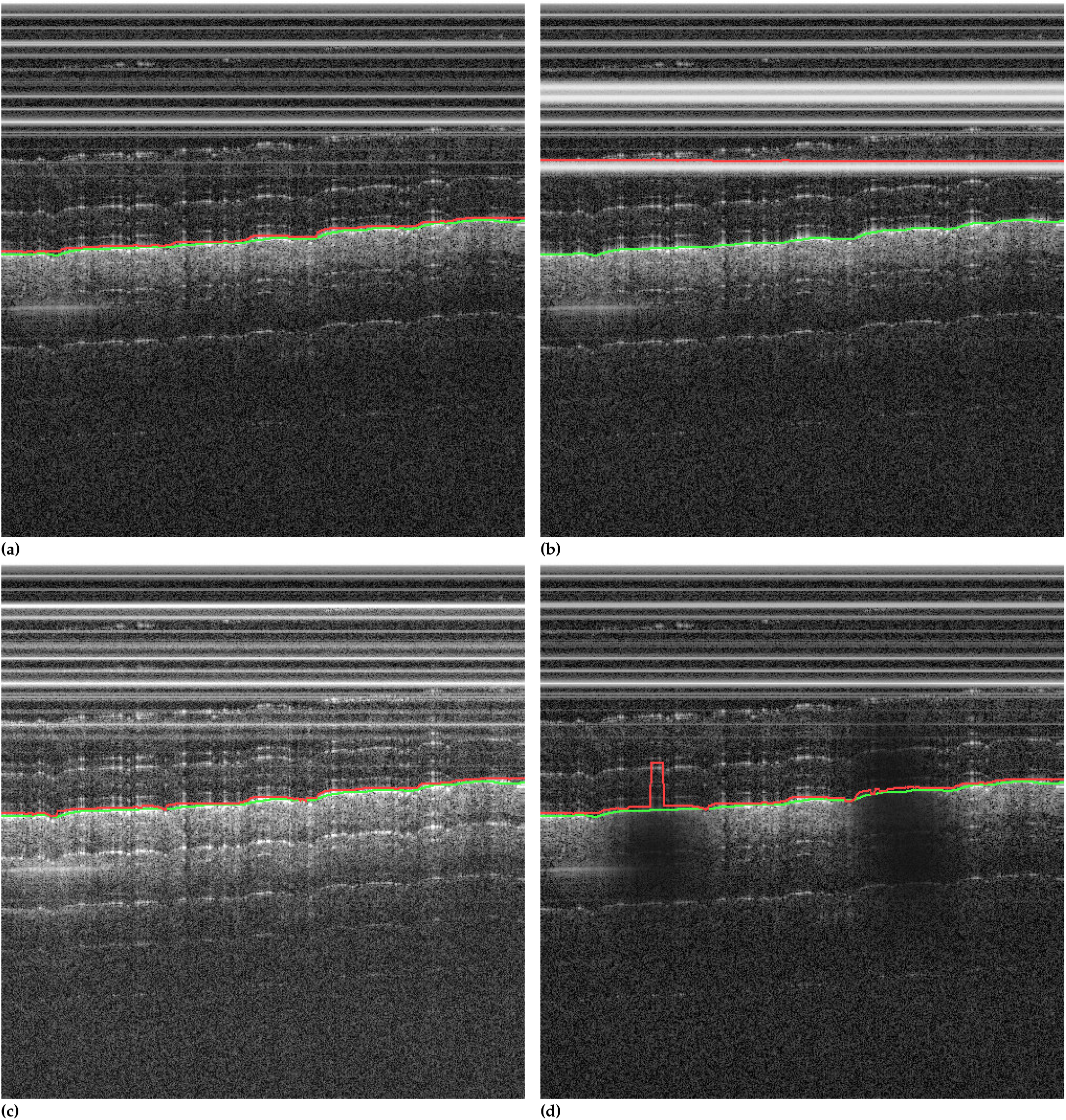
Representative examples of image corruptions caused by different artifacts used to evaluate robustness. (a) Clean OCT B-scan. (b) Horizontal stripe artifact, mimicking PM-fiber coherence stripe contamination. (c) Vertically shifted ghost copy, visible as a displaced duplicate structure that can create false surface predictions. (d) Local dropout/low contrast corruption, used as a broader surface visibility stress test. Operator definitions and parameter settings are provided in Table 10 (see Appendix, Section B). Green lines denote manual labels, and red lines denote classical method predictions.

## 3. RESULTS

### A. Main localization performance

Table 5 summarizes the clean test localization performance. All DL methods substantially reduced the clean set MAE relative to gradient-based baselines. Among the learned methods, the compact supervised baselines (FCBR and UNet) achieved the lowest clean set MAE and highest Acc@2px, while DINOCT traded roughly 0.5 px of clean MAE relative to the compact supervised baselines in exchange for a near-zero spike rate, which is especially important for autofocusing stability.

**Table 5.** Skin surface localization performance. Performance values are mean ±standard deviation across evaluation recordings; values for learned methods were averaged across five training seeds within each recording. Runtime is mean ±standard deviation over repeated per-B-scan timing measurements. Additional baseline implementation and architecture details are provided in Appendix, Section A.

| Method | MAE (px) ↓ | Bias (px) $\rightarrow 0$ | Acc@2px ↑ | SpikeRate@$\kappa$ (%) ↓ | Runtime (ms/B-scan) ↓ |
| --- | --- | --- | --- | --- | --- |
| GF | 37.174 ± 19.063 | -11.582 ± 14.146 | 0.420 ± 0.200 | 29.38 ± 7.62 | **2.131** ± 0.033 |
| GRAD-ENG | 18.252 ± 18.238 | -6.102 ± 18.145 | 0.542 ± 0.187 | 0.83 ± 0.42 | 20.051 ± 0.154 |
| TUNED-SOBEL-DC | 6.009 ± 7.767 | -4.226 ± 8.110 | 0.312 ± 0.288 | 1.80 ± 1.38 | 225.871 ± 3.106 |
| FCBR | 1.151 ± 2.203 | 0.408 ± 1.715 | 0.946 ± 0.069 | 0.42 ± 1.92 | 28.12 ± 0.42 |
| UNet | **1.108** ± 2.002 | 0.190 ± 1.050 | **0.949** ± 0.058 | 0.27 ± 1.19 | 26.02 ± 0.49 |
| DINOCT | 1.634 ± 2.345 | **-0.048** ± 1.663 | 0.821 ± 0.142 | **0.00** ± 0.01 | 34.15 ± 0.60 |

Runtime values reflect the deployment implementation on the evaluation computer; DINOCT was slower than the smaller learned baselines but remained within tens of milliseconds per B-scan. These results show that, under favorable imaging conditions, surface localization can be performed accurately using relatively compact supervised networks. DINOCT is therefore most useful in acquisitions affected by artifacts, where robustness to false structures caused by PM-fiber artifacts is more important than minimizing clean image MAE alone. A visual representation of the tradeoff can be seen in Fig. 9.

**Fig. 9.**
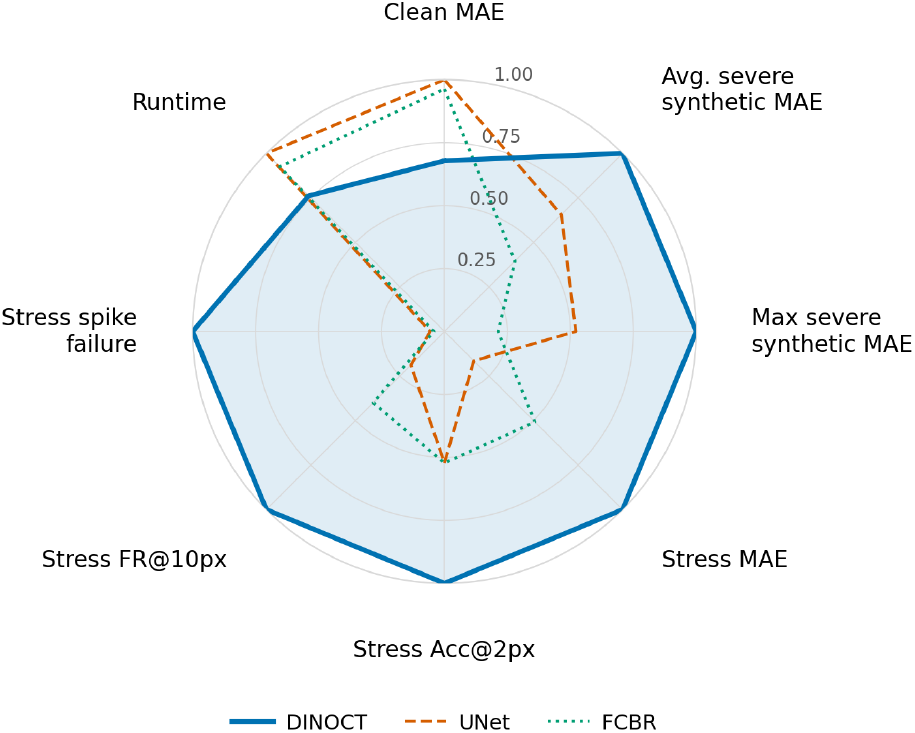
Relative performance of the learned methods. Each metric was normalized to the best result among the three methods. For the error, failure rate, and runtime, the best value was divided by the value for each method. For Acc@2px, the value for each method was divided by the best value. Larger radial values therefore indicate better relative performance. The original values are reported in Tables 5, 7, and 8. This figure illustrates the tradeoff between accuracy on clean images and robustness to synthetic artifacts and the stress recording. It is not an aggregate score. Four axes use results from the single stress recording.

### B. Artifact robustness

Table 6 summarizes MAE under severe synthetic corruption and on the stress recording. DINOCT achieved the lowest MAE under severe stripe and severe dropout corruption, while UNet achieved the lowest MAE under severe ghost corruption, where the three learned methods were closely matched (UNet 2.334, DINOCT 2.892, FCBR 2.931 px). On the stress recording, DINOCT achieved the lowest MAE among all evaluated methods. Full medium- and severe-corruption MAE and Acc@2px results are provided in Tables 15 and 16 (see Appendix, Section E).

**Table 6.**
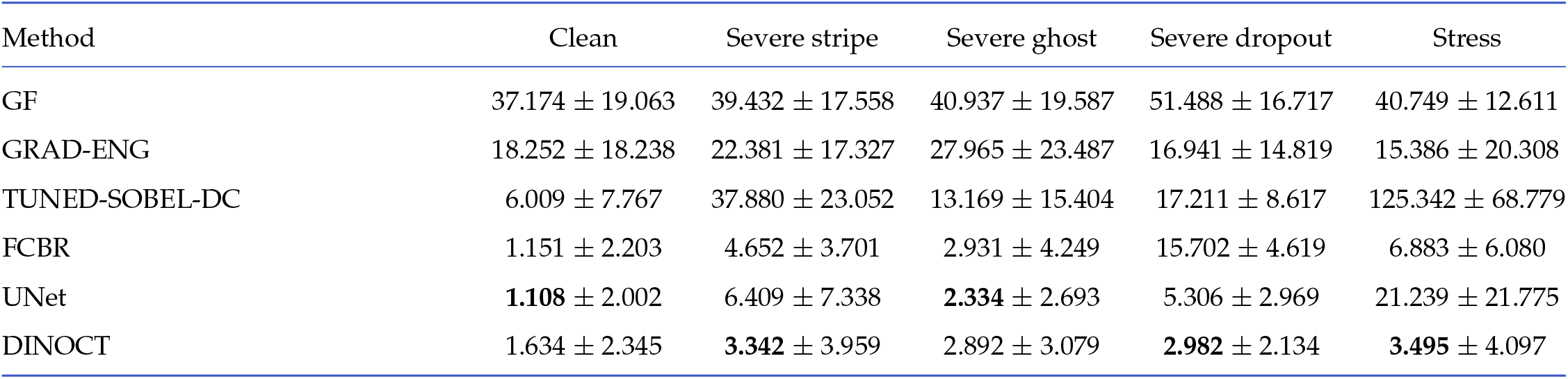
Robustness performance, MAE (px)↓. Clean and synthetic values are mean ±standard deviation across evaluation recordings; values for learned methods were averaged across five training seeds within each recording. The stress recording consists of one acquisition and is summarized across B-scans. Full medium- and severe-corruption results are provided in Appendix, Section E.

| Method | Clean | Severe stripe | Severe ghost | Severe dropout | Stress |
| --- | --- | --- | --- | --- | --- |
| GF | 37.174 ± 19.063 | 39.432 ± 17.558 | 40.937 ± 19.587 | 51.488 ± 16.717 | 40.749 ± 12.611 |
| GRAD-ENG | 18.252 ± 18.238 | 22.381 ± 17.327 | 27.965 ± 23.487 | 16.941 ± 14.819 | 15.386 ± 20.308 |
| TUNED-SOBEL-DC | 6.009 ± 7.767 | 37.880 ± 23.052 | 13.169 ± 15.404 | 17.211 ± 8.617 | 125.342 ± 68.779 |
| FCBR | 1.151 ± 2.203 | 4.652 ± 3.701 | 2.931 ± 4.249 | 15.702 ± 4.619 | 6.883 ± 6.080 |
| UNet | **1.108** ± 2.002 | 6.409 ± 7.338 | **2.334** ± 2.693 | 5.306 ± 2.969 | 21.239 ± 21.775 |
| DINOCT | 1.634 ± 2.345 | **3.342** ± 3.959 | 2.892 ± 3.079 | **2.982** ± 2.134 | **3.495** ± 4.097 |

**Table 7.**
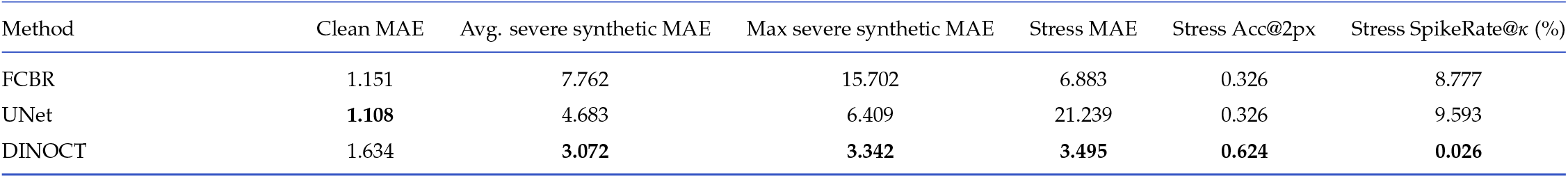
Aggregate robustness summary for DL models. The values for severe synthetic corruption are derived from the condition-level aggregate MAEs reported in Table 6. Average severe synthetic MAE is the arithmetic mean across the severe stripe, ghost, and dropout conditions; maximum severe synthetic MAE is the largest of those three values. The stress recording consists of one acquisition and is summarized across B-scans.

**Table 8.** Catastrophic failure rates for B-scans in the stress recording (%). Rates for learned methods average the per-seed failure indicator across five training seeds (the lower the better).

| Method | FailureRate@5px | FailureRate@10px | B-scan spike-failure rate |
| --- | --- | --- | --- |
| GF | 100.000 | 100.000 | 100.000 |
| GRAD-ENG | 94.040 | 29.801 | 99.338 |
| TUNED-SOBEL-DC | 99.338 | 96.689 | 100.000 |
| FCBR | 42.119 | 20.530 | 97.351 |
| UNet | 60.795 | 43.576 | 72.185 |
| DINOCT | **20.132** | **8.212** | **4.106** |

Table 7 compares robustness across artifact conditions among the methods evaluated in this study. Although the compact supervised baselines achieved the lowest clean MAE, DINOCT achieved the lowest average MAE across the severe synthetic conditions, the lowest maximum MAE across those conditions, and the lowest MAE, highest Acc@2px, and lowest spike rate on the stress recording among the learned methods.

Relative to clean MAE, the average severe synthetic MAE increased by 6.74× for FCBR, 4.23× for UNet, and 1.88× for DINOCT. On the stress recording, MAE increased by 5.98× for FCBR, 19.17× for UNet, and 2.14× for DINOCT.

These ratios should be interpreted cautiously because clean set errors differ across models, but they reinforce the trend that DINOCT degraded less under different artifact stresses.

Table 8 reports catastrophic failure rates for B-scans in the stress recording. DINOCT reduced FailureRate@10px from 20.5% for FCBR and 43.6% for UNet to 8.2%, and reduced the B-scan spike-failure rate from 97.4% (FCBR) and 72.2% (UNet) to 4.1%. Thus, the robustness advantage of DINOCT is best interpreted as fewer large errors and greater stability in conditions with artifacts, rather than uniformly better accuracy on clean scans.

### C. Qualitative examples

Figure 10 compares the strongest classical baselines and learned methods on the same OCT images. The top four rows show clean test examples, and the bottom two rows show examples from the stress recording. On the stress recording, DINOCT more consistently follows the visible surface and avoids large jumps. Additional failure cases of the classical method and further comparisons of all methods on the stress recording are provided in Appendix, Section F (Figs. 13 and 14).

**Fig. 10.**
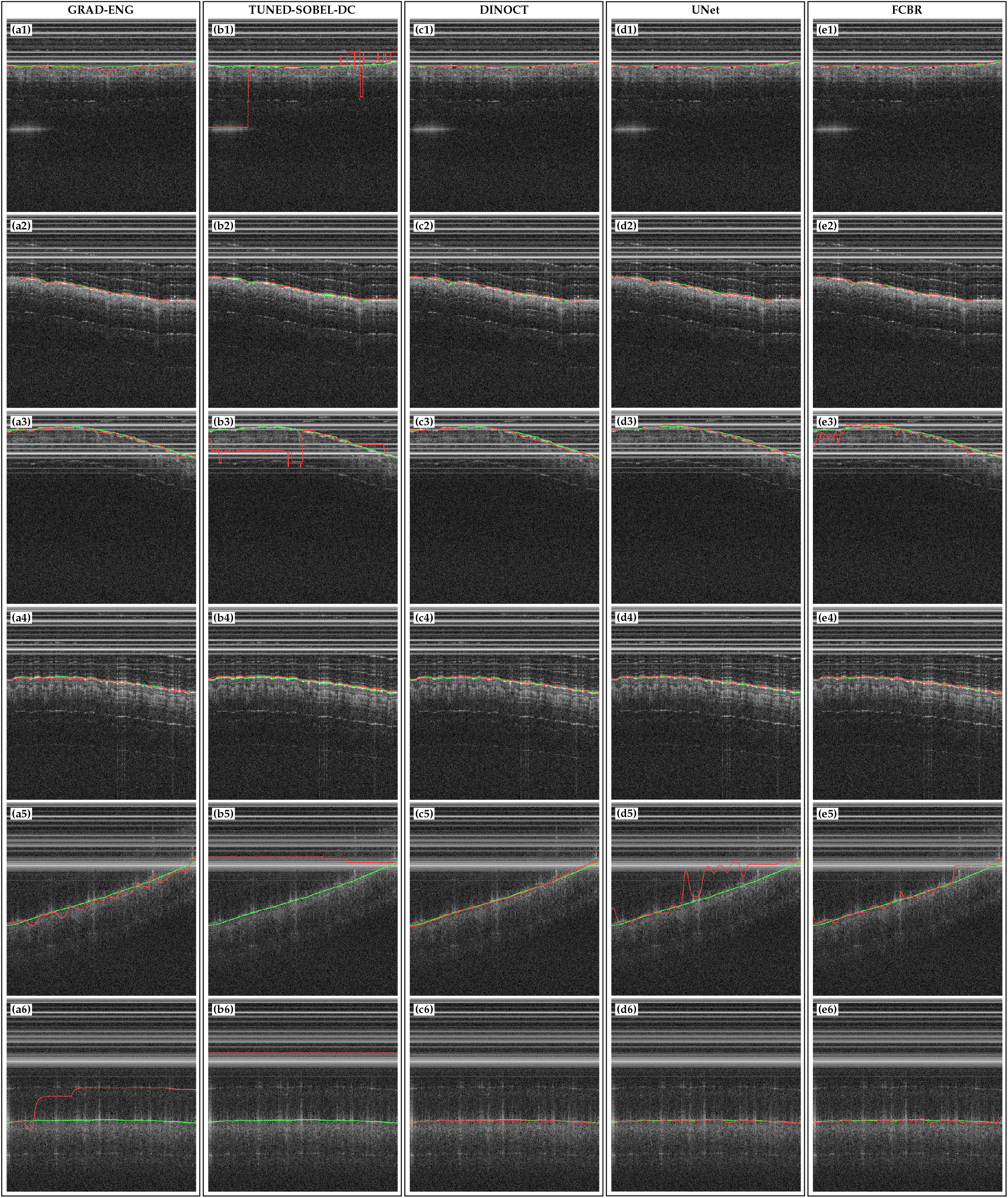
Qualitative comparison of results for surface localization obtained with different methods. Panels are indexed by method column and example row: (a1)–(a6) show **GRAD-ENG**, (b1)–(b6) show **TUNED-SOBEL-DC**, (c1)–(c6) show **DINOCT**, (d1)–(d6) show **UNet**, and (e1)–(e6) show **FCBR**. Rows 1–4 are clean test examples, and rows 5–6 are examples from the stress recording. Green lines denote manual labels, and red lines denote method predictions. Additional examples with real artifacts and classical failure cases are provided in Appendix, Section F.

### D. Examples of skin surface alignment

A representative OCT-guided alignment sequence is shown in Fig. 11; the closed loop positioning accuracy was not quantified 752 in this study.

**Fig. 11.**
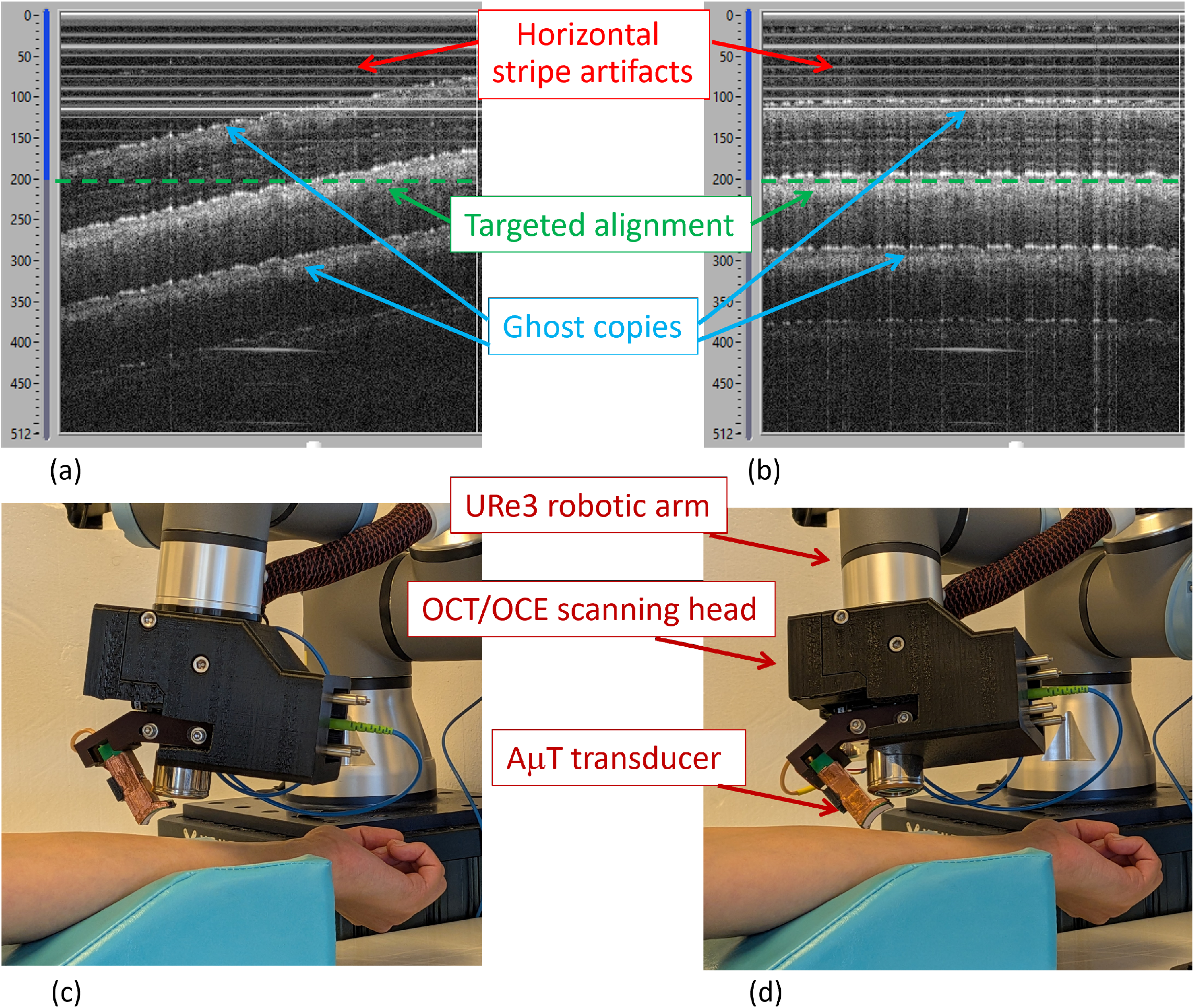
Representative alignment using OCT guidance. (a,b) Structural OCT preview images before and after alignment. Red arrows indicate horizontal stripe artifacts, blue arrows indicate ghost copies, and the dashed green line marks the target surface depth. (c,d) Corresponding photographs of the OCT/OCE head on the robot before and after alignment. This example illustrates the alignment procedure; positioning accuracy was not quantified in this study.

## 4. DISCUSSION

On clean test scans, the compact supervised baselines achieved the lowest MAE and highest accuracy at the tolerance of about 2 px, whereas DINOCT maintained competitive accuracy with a near-zero spike rate. DINOCT showed its strongest advantage under artifact stress: among the learned methods, it achieved the lowest average MAE across the severe synthetic conditions, the lowest maximum MAE across those conditions, and the best performance on the stress recording.

The severe synthetic conditions are unlikely to represent routine OCT imaging. Nevertheless, they show how the methods behave when image quality is substantially degraded. DINOCT maintained stable localization with practical accuracy and runtime. This tradeoff is relevant to robotic OCT/OCE because alignment errors can affect reconstructed mechanical properties [5]. Measurements in anisotropic skin must be repeated in several propagation directions, and the cylindrical transducer requires accurate head alignment to produce a narrow line source [4]. PM fibers provide the stable phase and polarization during robot motion but also produce structured image artifacts. DINOCT should therefore be understood as a method that improves localization stability in the presence of these artifacts, rather than as a method that maximizes accuracy on clean images.

For the present alignment task, mean errors of a few pixels were considered acceptable for OCE applications; suppressing rare large localization failures was more important than reducing mean pixel errors. For an order-of-magnitude comparison, a representative skin wave speed of approximately 4 m/s over the recorded 0.2–2 kHz frequency range corresponds to mechanical wavelengths of approximately 2–20 mm [1]. Using the axial calibration of 65 pixels/mm, a localization error of 3–4 pixels corresponds to approximately 0.05–0.06 mm. Because the calibration was approximate, all formal method comparisons remain reported in pixels.

Learned methods are compared as complete localization systems rather than methods isolating the effect of the SSL backbone. DINOCT combines self-supervised pretraining, adaptation with LoRA, optimization with SAM and EMA, and a constrained curve decoder. UNet and FCBR were trained from scratch using their respective supervised procedures.

The size of the present SSL pool is at or below the smallest regimes considered in most generic visual SSL studies. El-Nouby et al. included an approximately 8,000-image target dataset and found that masked or denoising objectives were more tolerant of limited data than DINO alone; they also observed degradation under excessively long pretraining schedules [35]. Konstantakos et al. defined low data pretraining as 50,000–300,000 images and reported strong dependence on the pretraining objective, architecture, downstream task, and domain [36]. Consequently, results from these classification studies do not predict performance for OCT surface localization, but they motivate checkpoint selection based on downstream tasks and controlled comparison of discriminative and masked/denoising pretraining objectives. Exploratory supervised label-efficiency results for the present dataset are reported in Appendix, Section C (Fig. 12 and Table 11).

**Fig. 12.**
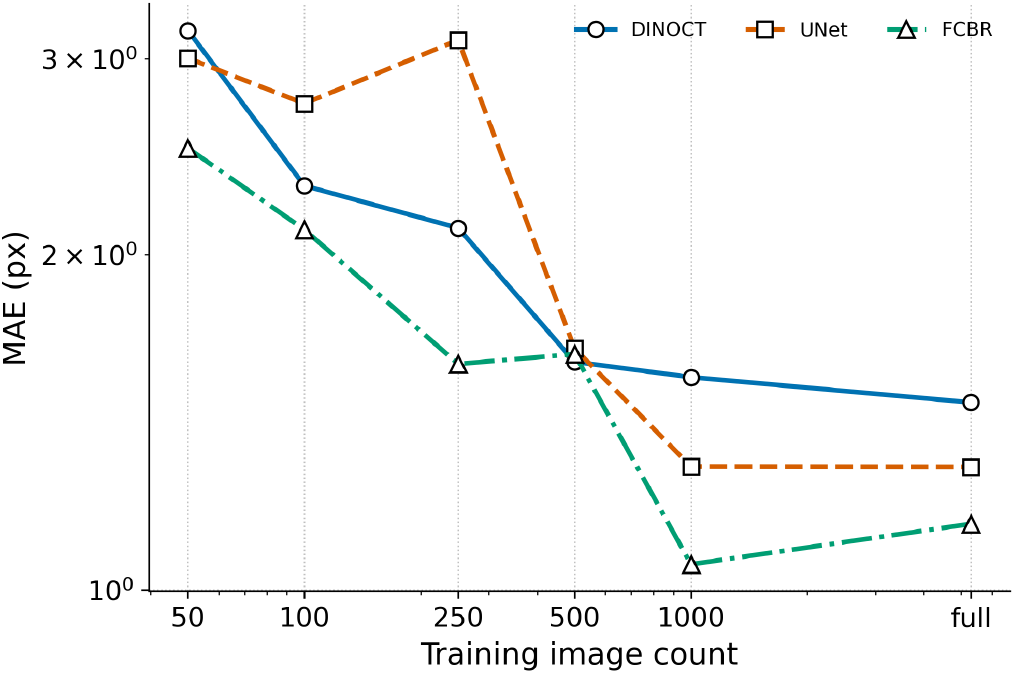
Clean test mean absolute error (MAE) versus labeled training B-scan count, shown with a logarithmic Y-axis. The corresponding full data multi-seed results are reported in Tables 5 and 6.

Backbone development showed that stable self-supervised optimization and robust downstream transfer were distinct objectives. Reducing the clustering prototype count improved early optimization consistency but weakened downstream transfer, while changes to KoLeo scheduling [41], auxiliary denoising reconstruction, and Gram-based feature consistency [42] did not yield reproducible robustness gains for the tested settings. Stronger augmentation showed no benefit at reduced training length but improved robustness when methods were compared at full training length. Denoising-based self-supervision has shown value for 3-D retinal OCT segmentation [37], but our experiment added denoising as an auxiliary term to DINO/iBOT and was not a reproduction of standalone denoising pretraining. The retained DINO/iBOT backbone continued to improve on frozen feature artifact localization after clean validation accuracy had plateaued. Therefore, checkpoint selection relied on downstream task probes rather than SSL loss or label free representation statistics alone.

The component sensitivity analysis in Appendix, Section D (Table 12) indicates that removing LoRA most consistently reduced robustness to real artifacts. Clean MAE was essentially unchanged, but MAE on the stress recording increased from 3.50 to 3.90 px, and Acc@2px decreased beyond the variation across seeds, with no meaningful improvement under any corruption. Freezing the backbone normalization parameters produced mixed effects: MAE improved under severe stripe and dropout corruption but worsened sharply under severe ghost corruption, and MAE on the stress recording also increased. Replacing SAM with AdamW slightly improved MAE on clean and synthetically corrupted images while increasing the error on real artifacts. EMA, curvature regularization, and the tested learning rate schedule produced changes within the variation across seeds.

This study has some limitations. First, all data were acquired on a single in-house platform, although there were minor changes in hardware configuration across recordings. Second, the stress recording contained only one acquisition in a small data regime. Its performance may therefore be sensitive to the composition of the training and validation split used during development. These results should be interpreted as evidence of robustness in challenging cases with artifacts, not as an estimate of performance across a broader out-of-distribution population. Third, the human subject data used in this study were obtained from healthy volunteers without known skin disease. Robustness in grafted tissue, burn scars, wounds, highly irregular skin surfaces, or other abnormal tissue structures was not evaluated. Therefore, the low observed failure rates under the current stress tests should not be interpreted as guaranteed performance across all clinically abnormal skin conditions. This subject will be a focus of our forthcoming studies and may require corresponding DINOCT algorithm tuning and additional network training. Finally, reference centerlines were manually annotated, but agreement between annotators was not measured. Differences near one pixel should therefore be interpreted in light of annotation uncertainty. DINOCT may also be useful for other OCT tasks that require reliable localization during image acquisition.

Many OCT workflows require fast and stable localization of interfaces or other geometric features, such as tissue surfaces, retinal or corneal boundaries, lumen interfaces, or probe-related structures, rather than full pixel-wise semantic segmentation. For these problems, a self-supervised OCT backbone can be reused as a feature extractor and coupled to a simple task-specific prediction head. This division is attractive when manual annotations are limited or when imaging artifacts differ across system configurations. The present curve decoder was intentionally constrained for single surface localization and therefore is not a replacement for dense segmentation networks. However, the same backbone adaptation strategy could be combined with alternative decoders for multi-layer boundary detection, multi-segment surfaces, or dense segmentation when those outputs are required. The main practical limitation is the computational cost of SSL pretraining, together with the need to choose backbone resolution and crop settings that preserve small spatial features in OCT images. Once such a representation is available, down-stream adaptation can be comparatively lightweight. Future work could evaluate joint-embedding predictive architectures such as LeJEPA [52] as a potential lower cost and more stable alternative to DINO/iBOT SSL for learning OCT backbones.

Finally, the current model assumes a single continuous visible surface and does not explicitly represent sharp discontinuities such as cuts or wide interruptions. In such cases, the model may either interpolate across the discontinuity by producing a full *W*-column curve or assign a high no-curve probability to affected columns or frames, neither of which is ideal for discontinuity-aware analysis. Extending the method to support breakpoint detection or multi-segment surface representations is another important direction for future work.

## 5. CONCLUSION

We presented DINOCT, a robust OCT-based framework for skin surface localization in robotic A*µ*T-OCE in the presence of structured PM-fiber artifacts. On the clean test set, DINOCT achieved an MAE of 1.634 ± 2.345 px, Acc@2px of 0.821, and a near-zero spike rate. The compact supervised baselines achieved the lowest clean-set MAE (UNet 1.108 px) and highest Acc@2px (UNet 0.949), indicating that DINOCT is best understood as a stability-oriented localizer rather than a method for improving clean image accuracy. Under artifact stress, DINOCT achieved the lowest average MAE across severe synthetic conditions among DL methods, 3.072 px, and the lowest maximum MAE across those conditions, 3.342 px. On the stress recording, DINOCT achieved the lowest MAE, 3.495 px, the highest Acc@2px, 0.624, and the lowest catastrophic failure rates. Specifically, DINOCT reduced FailureRate@10px to 8.2%, compared with 20.5% for FCBR and 43.6% for UNet, and reduced the B-scan spike-failure rate to 4.1%. DINOCT ran in 34.15 ± 0.60 ms per B-scan on the deployment computer, corresponding to approximately 29 B-scans/s. For a sparse OCT data volume (6 B-scans) used for surface fitting and OCT/OCE head alignment, this corresponds to approximately 0.2 s of DINOCT processing time for surface localization, excluding robot motion. These results establish DINOCT as a stable localizer for artifact-prone robotic OCT/OCE workflows. It is especially useful when reducing rare large localization failures is more important than further improving the already high accuracy on clean images.

## APPENDIX

### A. Implementation and baseline details

This section provides implementation details for the comparison methods summarized in Section H and supplements the results. The primary clean and robustness results are reported in Sections A and B. Exact implementations and configurations are provided in the accompanying code release [45].

#### GF-B

GF-B is a background-subtracted modification to the maximum gradient fitting method. GF-B first subtracts a reference background image from the input, applies row-wise DC suppression, and then uses the same gradient scoring and column-wise peak-picking procedure as GF.

#### GRAD-SG

GRAD-SG first applied a 5 × 5 median filter, followed by row-wise DC suppression, a 3 × 11 median filter, and Gaussian smoothing with *σ* = 3. The surface was selected from the column-wise maximum of the vertical gradient and then processed using a scalar recursive filter (*q* = 0.01, *r* = 0.5), local outlier correction, and a Savitzky–Golay filter with window length 15 and polynomial order 3.

#### A.1 Additional learned baseline details

The main paper summarizes the U-Net and FCBR-style baselines in Section H. The learned baselines were trained from scratch on the same input resolution and evaluated on the same labeled train/validation/test split used for DINOCT post-training. All methods were evaluated using the same manually annotated centerlines and the same pixel domain metrics. See Table 9 for architecture summaries.

**Table 9.** Additional learned baseline architectures and parameter counts for the UNet and FCBR-style methods summarized in Section H.

| Model | Architecture summary | Parameters |
| --- | --- | --- |
| UNet | 4-level encoder–decoder with skip connections; channels 32/64/128/192; bottleneck 256; curve-distribution head for per-column surface localization | 3,689,954 |
| FCBR-style | Shallow fully convolutional boundary regressor with $5 \times 5$ stem, depthwise separable downsampling blocks, 5-block dilated context stack, and fusion block | 131,234 |

To ensure a fair comparison, all learned methods were trained using a matched sample budget of approximately 192,000 supervised samples: DINOCT used 1,500 steps at batch size 128, the FCBR-style baseline 6,000 steps at batch size 32, and the UNet baseline 16,000 steps at batch size 12. Under substantially longer training, the from-scratch baselines began to overfit the clean training distribution, which further widened their real artifact stress gap relative to DINOCT.

##### UNet

The UNet baseline was implemented as a four-level encoder–decoder with skip connections. Each encoder and decoder block used two 3 × 3 convolutions with GroupNorm and GELU activation. The encoder stages used 32, 64, 128, and 192 channels, followed by a 256-channel bottleneck. The decoder used bilinear upsampling and feature concatenation with the corresponding encoder outputs. The final feature map was passed to a curve-distribution head that predicted *H* depth logits and one no-curve logit for each image column. The implemented model contained 3,689,954 trainable parameters.

##### FCBR-style boundary regressor

The FCBR-style baseline was implemented as a lightweight fully convolutional boundary regressor. The model used a 5 × 5 convolutional stem with stride (2, 1), followed by depthwise separable convolutional blocks with channel dimensions 32, 64, and 96. After the downsampling stages, a 5-block dilated context stack with dilation sequence 1– 2–4–2–1 was applied, followed by a fusion module consisting of a 1 × 1 convolution and a 3 × 3 convolution. The fused features were passed to a curve-distribution head that predicted *H* depth logits and one no-curve logit for each image column. The total parameter count of the implemented model was 131,234, all of which were trainable. We report this baseline as an FCBR-style reimplementation rather than an exact historical reproduction.

##### Learned-baseline training

Both learned baselines were trained from scratch using the curve-distribution objective described in Section C. The loss used a Gaussian depth target with *σ*_*z*_ = 1.5 px, *λ*_pred_ = 1.0, *λ*_smooth_ = 0.05, a non-visible sample weight of 5.0, and *ϵ*none = 0.02. Optimization used AdamW with an initial learning rate of 10^™3^, weight decay 5 × 10^™4^, 50 warm-up steps, and cosine decay to 0.1 of the initial learning rate. Inputs were resized to 512 × 500, converted to three channels, and standardized independently for each image. No data augmentation was used. The checkpoint with the lowest validation MAE was retained.

## B. Synthetic corruption operators

Synthetic corruptions were applied independently to each clean test B-scan at evaluation time. Let *I*(*r, x*) denote the input OCT image intensity at axial pixel row *r* and lateral column *x*, and let *Ī* denote the image mean. The operators below were used to generate medium and severe corruption conditions. The main robustness protocol is described in Section J, and representative examples are shown in Fig. 8.

### Stripe corruption

Stripe corruption added bright horizontal coherence-like bands (see Section J and Fig. 8(b)). For each stripe *m*, a row center *c*_*m*_ was sampled near the upper part of the image with small jitter, and a vertical Gaussian band profile

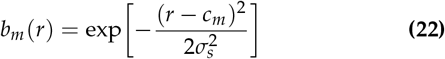

was used to blend the image toward a bright target intensity *T*_*s*_:

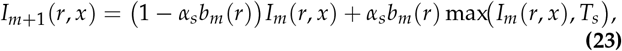

where *α*_*s*_ is the stripe opacity and the bandwidth is set from the nominal thickness *t* as *σ*_*s*_ = *t*/2.2 px. The values of parameters are listed in Table 10.

### Ghost corruption

Ghost corruption added a vertically shifted copy of the image (see Section J and Fig. 8(c)), defined as

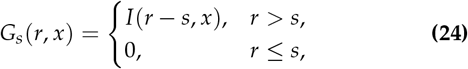

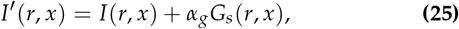

where *s* is the axial shift and *α*_*g*_ is the ghost opacity. The values of parameters are listed in Table 10.

### Dropout and low contrast corruptions

Dropout/low contrast corruptions were simulated by localized regions with reduced intensity and contrast using separable raised-cosine masks (see Section J and Fig. 8(d)). First, a contrast compressed and intensity scaled image was formed:

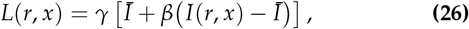

where *β* is the contrast scale and *γ* is the intensity scale. A separable raised cosine mask *M*(*r, x*) ∈ [0, 1] was then used to blend the original image with *L*:

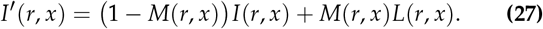

The mask parameters are listed in Table 10.

**Table 10.** Synthetic corruption parameters used for robustness evaluation. These settings correspond to the operators described in Section J and illustrated in Fig. 8. Dropout region size denotes the nominal full raised-cosine mask width × height; the implementation applies ±15% random size variation.

| Artifact | Parameter | Medium | Severe |
| --- | --- | --- | --- |
| Stripe | Number of bands | 2 | 3 |
|  | Nominal thickness | 6 px | 9 px |
| | Opacity $\alpha_s$ | 0.78 | 0.92 |
| | Target intensity $T_s$ | 236 | 248 |
|  | Upper sampling band | 200 px | 200 px |
| Ghost | Shift $s$ | 24 px | 40 px |
| | Opacity $\alpha_g$ | 0.25 | 0.45 |
| Dropout | Number of regions | 2 | 3 |
| | Nominal region size | 112 × 160 px | 160 × 220 px |
| | Contrast scale $\beta$ | 0.46 | 0.30 |
| | Intensity scale $\gamma$ | 0.58 | 0.42 |

## C. Additional supervised label efficiency results

The full-data DINOCT architecture and training objective are described in Sections B and C, with the performance summary reported in Sections A and B.

We also performed an exploratory analysis (as described below) using one seed and reduced numbers of supervised labeled images to assess how the complete DINOCT recipe behaved as the manual annotation volume decreased. This analysis evaluated supervised label efficiency rather than total data efficiency because DINOCT used the same unlabeled SSL pre-training pool, while the number of labeled visible B-scans used for post-training was varied.

On the clean test set, DINOCT achieved the lowest MAE at the 500-label budget; FCBR achieved the lowest MAE at the remaining reported budgets. On the stress recording, DINOCT achieved the lowest MAE among the learned methods at every label budget, by a large margin at small budgets (e.g. 5.01 vs 17.6–21.6 px at 50 labels), and maintained a near zero spike rate, although its MAE for real artifacts was not strictly monotonic with label count.

These results from one seed should be interpreted as label-budget trends for the complete training recipes rather than as variance-controlled comparisons or an isolation of the effects of SSL pretraining and LoRA adaptation. This behavior is consistent with broader reports that in-domain SSL can be advantageous for specialized imaging domains [36]; however, the present comparison does not isolate SSL pretraining from the downstream architecture and adaptation strategy.

**Table 11.** Exploratory single-seed supervised label-efficiency summary, MAE (px)↓. The “full” row corresponds to the same single seed and may differ from the multi-seed results in Tables 5 and 6.

| Label budget | Clean test MAE |  |  | Stress MAE |  |  |
| --- | --- | --- | --- | --- | --- | --- |
|  | DINOCT | UNet | FCBR | DINOCT | UNet | FCBR |
| 50 | 3.179 | 3.002 | **2.492** | **5.009** | 21.597 | 17.561 |
| 100 | 2.306 | 2.733 | **2.106** | **5.452** | 21.801 | 17.802 |
| 250 | 2.113 | 3.117 | **1.595** | **5.151** | 26.941 | 11.750 |
| 500 | **1.602** | 1.647 | 1.628 | **4.032** | 21.147 | 12.117 |
| 1000 | 1.552 | 1.290 | **1.054** | **3.686** | 10.505 | 9.609 |
| Full | 1.473 | 1.289 | **1.146** | **3.082** | 26.069 | 7.188 |

## D. Component sensitivity

The reference DINOCT configuration is described in Section B, Fig. 3, and Table 2; the main robustness results are reported in Section B. To assess which parts of the DINOCT training configuration contribute to robustness, we performed a single component sensitivity analysis on a controlled reference training run (Table 12). This analysis uses the same five seeds as the main evaluation; values are aggregated under the sensitivity-analysis protocol (mean ± std across seeds). Results of the analysis support the same clean accuracy versus robustness tradeoff observed in Sections A and B. DINOCT’s MAE on clean images remained below 2 px, which was adequate for the applications discussed here. Removing LoRA produced the clearest reduction in robustness to real artifacts. Clean MAE was essentially unchanged, whereas MAE on the stress recording increased from 3.50 to 3.90 px and Acc@2px decreased from 0.624 to 0.588.

Freezing the backbone normalization parameters produced mixed effects. MAE improved under severe stripe and dropout corruption, whereas MAE under severe ghost corruption worsened sharply (3.97 versus 2.89 px), and MAE on the stress recording increased by a margin similar to LoRA removal (3.90 versus 3.50 px). Restricting the norm adaptation degraded robustness on the stress recording while leaving in-distribution accuracy essentially unchanged (Table 13): freezing all backbone norms or adapting only the final norm changed clean MAE by at most 0.05 px but increased MAE on the stress recording by 0.40–0.62 px, an effect comparable to removing LoRA (0.40 px); adapting only the final norm did not recover this gap. We therefore keep all backbone norms trainable.

**Table 12.** Single component sensitivity analysis relative to the reference DINOCT training and adaptation configuration. Clean and corruption columns report MAE (px), shown as mean ± standard deviation across five seeds ↓; the columns for the stress recording report MAE (px) ↓ and Acc@2px ↑. The reference configuration is summarized in Fig. 3 and Table 2 in the main text.

| Configuration | Change from Full | Clean | Severe stripe | Severe ghost | Severe dropout | Stress recording |  |
| --- | --- | --- | --- | --- | --- | --- | --- |
| | | | | | | MAE (px) ↓ | Acc@2px ↑ |
| Full | Reference configuration; all backbone norms trainable | 1.634 ± 0.224 | 3.342 ± 0.238 | 2.891 ± 0.153 | 2.982 ± 0.350 | 3.495 ± 0.266 | 0.624 ± 0.010 |
| -EMA | EMA = 0 | 1.596 ± 0.122 | 3.337 ± 0.275 | 3.080 ± 0.196 | 2.948 ± 0.322 | **3.475** ± 0.272 | 0.620 ± 0.014 |
| -Curvature | $\lambda_{\text{smooth}} = 0$ | 1.634 ± 0.224 | 3.342 ± 0.238 | 2.891 ± 0.153 | 2.982 ± 0.350 | 3.495 ± 0.266 | 0.624 ± 0.010 |
| -LoRA | LoRA = 0 | **1.529** ± 0.119 | 3.554 ± 0.272 | 2.841 ± 0.276 | 3.139 ± 0.146 | 3.899 ± 0.188 | 0.588 ± 0.016 |
| AdamW | SAM → AdamW | 1.566 ± 0.082 | 3.292 ± 0.148 | **2.530** ± 0.191 | 2.951 ± 0.157 | 3.606 ± 0.241 | 0.619 ± 0.018 |
| +LR schedule | lr_warmup=10, min_lr_mult=0.95 | 1.598 ± 0.193 | 3.333 ± 0.254 | 2.821 ± 0.177 | 3.034 ± 0.325 | 3.494 ± 0.271 | **0.625** ± 0.009 |
| -BackboneNorms | all backbone norms → frozen | 1.606 ± 0.027 | **2.850** ± 0.215 | 3.971 ± 0.240 | **2.208** ± 0.086 | 3.898 ± 0.220 | 0.594 ± 0.022 |

The optimizer choice could modify the relationship between the accuracy for clean images versus robustness to real artifacts: AdamW slightly improved MAE for clean images and images with synthetic corruptions but increased the error for real artifacts. Removing EMA and/or the curvature term, or adding the tested learning rate schedule produced changes within the variation across seeds. These results indicate that LoRA removal produced the clearest degradation in robustness for the stress recording, whereas the optimizer and choices for backbone normalization produced mixed effects on accuracy and robustness. EMA, curvature regularization, and the tested learning-rate schedule had relatively minor effects.

Table 14 reports a LoRA placement sensitivity analysis. Recent post-training experiments on language models reported benefits from broad LoRA placement, particularly in MLP layers [53]. Motivated by this observation, we evaluated LoRA coverage in the pointwise MLP layers of ConvNeXt. Under the present OCT post-training recipe, placing LoRA on the final six blocks gave the best average MAE, best maximum MAE across the severe corruptions and the highest accuracy for real artifacts (Acc@2px). Extending LoRA to all eighteen blocks worsened the maximum MAE across the severe corruptions and Acc@2px.

**Table 13.** Backbone normalization adaptation sensitivity analysis across five seeds. For each seed, average severe synthetic MAE was computed as the mean across the severe stripe, ghost, and dropout conditions, and maximum severe synthetic MAE was the largest of those three values. The table reports the mean ± population standard deviation of these per-seed summaries. The all-norms and frozen rows share their underlying runs with the Full and ™BackboneNorms configurations of Table 12.

| Configuration | Change | Clean MAE | Avg. severe synthetic MAE | Max severe synthetic MAE | Stress MAE | Stress Acc@2px |
| --- | --- | --- | --- | --- | --- | --- |
| All backbone norms | Reference | 1.634 ± 0.224 | 3.072 ± 0.066 | **3.479** ± 0.104 | **3.495** ± 0.266 | **0.624** ± 0.010 |
| Final norm only | all backbone norms → final norm only | **1.582** ± 0.031 | **2.975** ± 0.051 | 3.698 ± 0.350 | 4.119 ± 0.455 | 0.583 ± 0.032 |
| Frozen backbone norms | all backbone norms → frozen | 1.606 ± 0.027 | 3.010 ± 0.073 | 3.971 ± 0.240 | 3.898 ± 0.220 | 0.594 ± 0.022 |

**Table 14.** LoRA placement sensitivity analysis. Values are mean ± population standard deviation across five seeds. For each seed, average severe synthetic MAE was computed as the mean across the severe stripe, ghost, and dropout conditions, and maximum severe synthetic MAE was the largest of those three values. The reported values then average these per-seed summaries across seeds. Table 7 instead reports the maximum of the condition means after averaging across seeds. All variants keep the remaining training recipe fixed; the selected last-six-block placement is the main text configuration in Section B and Table 2.

| Configuration | Change | Clean MAE | Avg. severe synthetic MAE | Max severe synthetic MAE | Stress MAE | Stress Acc@2px |
| --- | --- | --- | --- | --- | --- | --- |
| No LoRA | LoRA = 0 | **1.529** ± 0.119 | 3.178 ± 0.083 | 3.600 ± 0.219 | 3.899 ± 0.188 | 0.588 ± 0.016 |
| Last 1 block | LoRA on final ConvNeXt block | 1.637 ± 0.138 | 3.211 ± 0.159 | **3.517** ± 0.223 | 3.941 ± 0.260 | 0.589 ± 0.014 |
| Last 3 blocks | LoRA on final 3 ConvNeXt blocks | 1.753 ± 0.256 | 3.318 ± 0.085 | 3.640 ± 0.103 | 4.020 ± 0.244 | 0.590 ± 0.015 |
| Last 6 blocks | Prior default placement | 1.634 ± 0.224 | **3.072** ± 0.066 | **3.479** ± 0.103 | 3.495 ± 0.266 | **0.624** ± 0.010 |
| Last 12 blocks | LoRA on final 12 ConvNeXt blocks | 1.699 ± 0.051 | 3.142 ± 0.091 | 3.515 ± 0.123 | 3.344 ± 0.102 | 0.614 ± 0.008 |
| All 18 blocks | LoRA on all ConvNeXt blocks | 1.703 ± 0.058 | 3.272 ± 0.105 | 3.898 ± 0.315 | **3.306** ± 0.273 | 0.576 ± 0.033 |

Adaptation capacity interacted with the label budget: deeper adaptation (all eighteen blocks) reduced error most clearly in the regime with few labels, whereas the moderate configuration was preferred at the full label budget (observations from one seed). Increasing spatial adaptation capacity by unfreezing the backbone depthwise convolutions improved clean accuracy but sharply degraded performance on real artifacts. This pattern of overfitting to clean images is consistent with the broader observation that robustness and clean-image accuracy can be distinct and sometimes competing objectives [54]. We deliberately did not augment post-training with the synthetic corruption operators used for evaluation, since doing so would train on the evaluation distribution. The reported robustness therefore reflects the features learned during pretraining and the subsequent adaptation rather than direct training on those synthetic artifacts.

## E. Full robustness tables

Tables 15 and 16 provide the full medium- and severe-corruption results underlying the summary in Section B and Tables 6 and 7. UNet achieved the best clean MAE and the lowest MAE under medium corruption and severe ghost corruption. DINOCT achieved the lowest MAE under severe stripe and dropout corruption and on the stress recording. Acc@2px showed a similar tradeoff: the compact supervised baselines performed best on clean images and synthetic corruptions at a tight tolerance, while DINOCT achieved the highest Acc@2px on the stress recording.

**Table 15.** Artifact robustness performance, MAE (px)↓. Clean and synthetic-corruption values are mean ±standard deviation across evaluation recordings; learned-method metrics were averaged across five training seeds within each recording. Classical methods are deterministic single runs. The stress recording contains one acquisition and is summarized across B-scans. These data expand the main text summary in Tables 6 and 7.

| Method | Clean | Medium stripe | Severe stripe | Medium ghost | Severe ghost | Medium dropout | Severe dropout | Stress |
| --- | --- | --- | --- | --- | --- | --- | --- | --- |
| GF | 37.174 ± 19.063 | 37.918 ± 18.612 | 39.432 ± 17.558 | 42.526 ± 19.735 | 40.937 ± 19.587 | 40.727 ± 18.008 | 51.488 ± 16.717 | 40.749 ± 12.611 |
| GF-B | 42.262 ± 18.544 | 42.647 ± 17.795 | 44.622 ± 16.928 | 46.504 ± 18.702 | 44.555 ± 18.801 | 45.125 ± 17.101 | 56.516 ± 15.699 | 65.722 ± 19.613 |
| GRAD-SG | 15.635 ± 17.034 | 16.152 ± 16.661 | 17.320 ± 15.828 | 15.623 ± 16.160 | 13.813 ± 15.197 | 18.352 ± 15.555 | 25.141 ± 13.157 | 14.840 ± 10.960 |
| GRAD-ENG | 18.252 ± 18.238 | 19.753 ± 18.076 | 22.381 ± 17.327 | 25.529 ± 21.613 | 27.965 ± 23.487 | 18.474 ± 16.127 | 16.941 ± 14.819 | 15.386 ± 20.308 |
| TUNED-SOBEL-DC | 6.009 ± 7.767 | 24.868 ± 22.987 | 37.880 ± 23.052 | 6.461 ± 8.248 | 13.169 ± 15.404 | 8.903 ± 7.671 | 17.211 ± 8.617 | 125.342 ± 68.779 |
| FCBR | 1.151 ± 2.203 | 1.705 ± 2.389 | 4.652 ± 3.701 | 1.557 ± 3.269 | 2.931 ± 4.249 | 4.941 ± 2.712 | 15.702 ± 4.619 | 6.883 ± 6.080 |
| UNet | **1.108** ± 2.002 | **1.519** ± 2.567 | 6.409 ± 7.338 | **1.354** ± 2.029 | **2.334** ± 2.693 | **1.307** ± 1.946 | 5.306 ± 2.969 | 21.239 ± 21.775 |
| DINOCT | 1.634 ± 2.345 | 2.078 ± 2.929 | **3.342** ± 3.959 | 1.681 ± 2.339 | 2.892 ± 3.079 | 1.664 ± 2.044 | **2.982** ± 2.134 | **3.495** ± 4.097 |

**Table 16.** Artifact robustness performance, Acc@2px↑. Clean and synthetic-corruption values are mean ±standard deviation across evaluation recordings; learned-method metrics were averaged across five training seeds within each recording. Classical methods are deterministic single runs. The stress recording contains one acquisition and is summarized across B-scans. These data expand the main text summary in Tables 6 and 7.

| Method | Clean | Medium stripe | Severe stripe | Medium ghost | Severe ghost | Medium dropout | Severe dropout | Stress |
| --- | --- | --- | --- | --- | --- | --- | --- | --- |
| GF | 0.420 ± 0.200 | 0.413 ± 0.194 | 0.394 ± 0.181 | 0.359 ± 0.195 | 0.344 ± 0.202 | 0.398 ± 0.185 | 0.334 ± 0.157 | 0.220 ± 0.123 |
| GF-B | 0.381 ± 0.187 | 0.374 ± 0.182 | 0.357 ± 0.170 | 0.328 ± 0.182 | 0.314 ± 0.190 | 0.362 ± 0.174 | 0.306 ± 0.147 | 0.156 ± 0.087 |
| GRAD-SG | 0.361 ± 0.242 | 0.356 ± 0.235 | 0.340 ± 0.216 | 0.347 ± 0.240 | 0.372 ± 0.236 | 0.332 ± 0.216 | 0.291 ± 0.179 | 0.206 ± 0.114 |
| GRAD-ENG | 0.542 ± 0.187 | 0.516 ± 0.181 | 0.485 ± 0.179 | 0.446 ± 0.193 | 0.378 ± 0.211 | 0.514 ± 0.166 | 0.496 ± 0.161 | 0.234 ± 0.131 |
| TUNED-SOBEL-DC | 0.312 ± 0.288 | 0.198 ± 0.180 | 0.122 ± 0.123 | 0.291 ± 0.280 | 0.259 ± 0.249 | 0.309 ± 0.275 | 0.282 ± 0.247 | 0.015 ± 0.053 |
| FCBR | 0.946 ± 0.069 | 0.852 ± 0.100 | 0.640 ± 0.167 | 0.890 ± 0.114 | **0.759** ± 0.176 | 0.887 ± 0.064 | 0.731 ± 0.071 | 0.326 ± 0.166 |
| UNet | **0.949** ± 0.058 | **0.883** ± 0.077 | **0.685** ± 0.137 | **0.891** ± 0.119 | 0.738 ± 0.213 | **0.930** ± 0.061 | **0.799** ± 0.075 | 0.326 ± 0.165 |
| DINOCT | 0.821 ± 0.142 | 0.737 ± 0.131 | 0.583 ± 0.150 | 0.808 ± 0.121 | 0.519 ± 0.233 | 0.804 ± 0.142 | 0.728 ± 0.143 | **0.624** ± 0.287 |

## F. Additional qualitative examples

These examples supplement the main qualitative comparison in Section C and Fig. 10. Fig. 13 is included to illustrate the dominant failure mechanisms observed for classical gradient-based detectors: locking to ghost reflections, responding to horizontal stripe/DC artifacts, and responding to localized high-intensity features. These examples help explain the high MAE, spike rate, and failure rates for B-scans observed for the classical baselines in conditions with artifacts.

Figure 14 provides additional examples from the stress recording and compares all methods on the same B-scans. These examples complement the catastrophic failure rates in Table 8 by showing how DINOCT avoids large surface jumps in cases where methods trained from scratch or classical methods can track artifact structures.

**Fig. 13.**
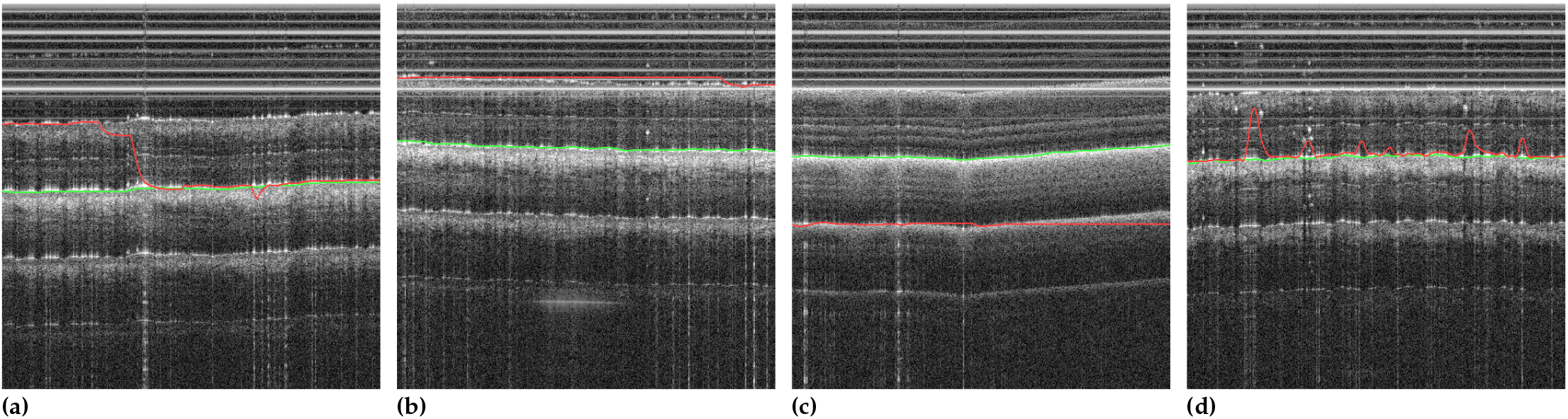
Representative failure modes for classical gradient-based methods. The green line denotes the manual reference centerline and the red line denotes the predicted centerline for the following examples: (a) Switching between a ghost reflection and the true surface, (b) Dominance of horizontal DC/stripe artifacts over the surface intensity response, (c) Detection of ghost reflections instead of the true surface, (d) Detection of localized high-intensity features associated with hair. These cases supplement the main text comparison in Fig. 10.

**Fig. 14.**
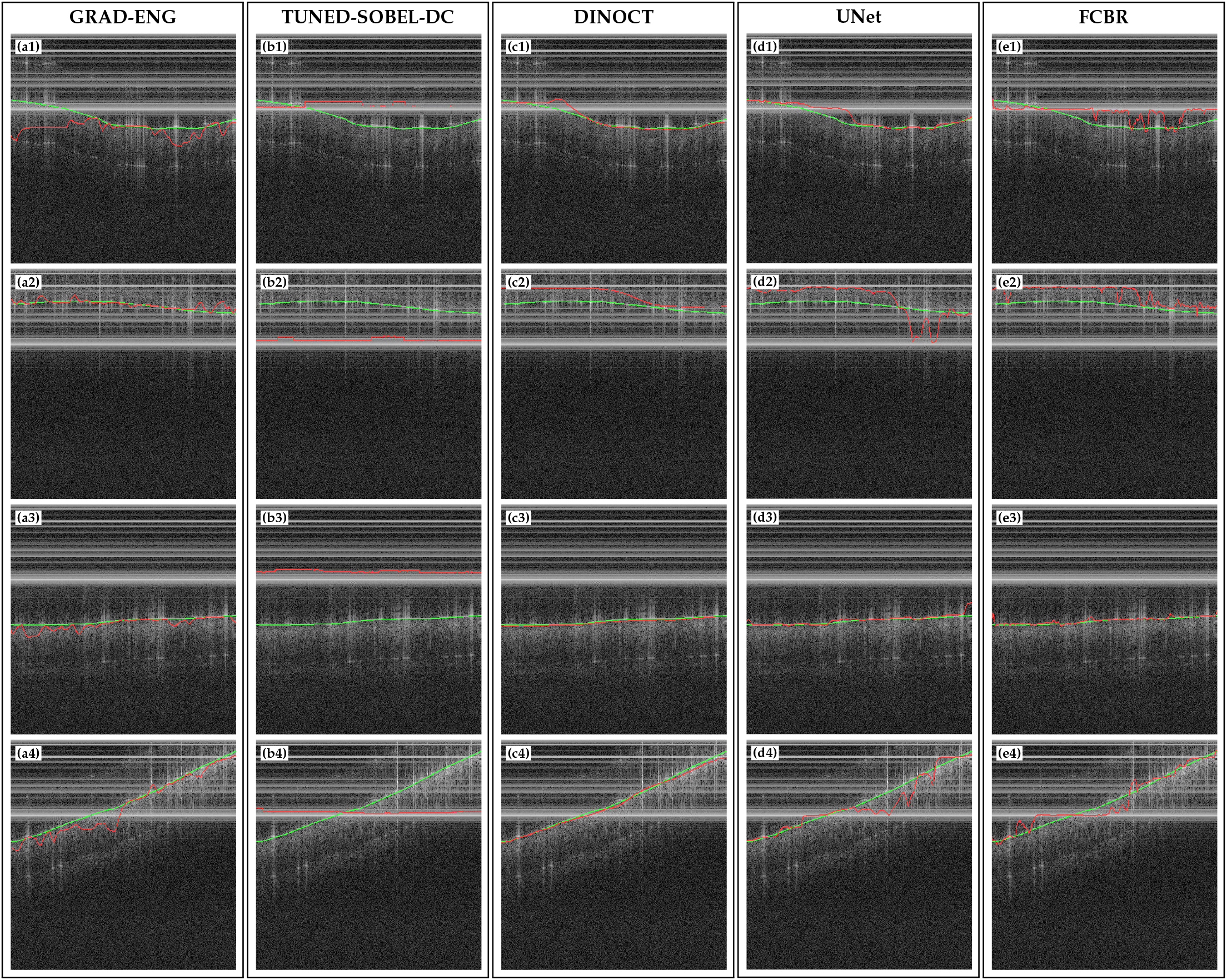
Additional qualitative examples from the stress recording. Panels are indexed by method column and example row: (a1)– (a4) show **GRAD-ENG**, (b1)–(b4) show **TUNED-SOBEL-DC**, (c1)–(c4) show **DINOCT**, (d1)–(d4) show **UNet**, and (e1)–(e4) show **FCBR**. Each row shows the same OCT B-scan evaluated by all methods. Green lines denote manual labels, and red lines denote method predictions. These examples extend main text Fig. 10 and complement the catastrophic failure rates in Table 8.

## Acknowledgment

This work was supported, in part, by NIH grants R01EY035647 and R01AR077560 and the Department of Bioengineering at the University of Washington.

## Data Availability

The OCT dataset and evaluation scripts underlying this work are available at [45, 55]. The released dataset includes the structural OCT images and split/evaluation resources used for surface-localization experiments, subject to dataset license and access terms. Additional robot platform code and workflow videos demonstrating autofocus and full scan operation are available at [56].

